# Direct evidence for transport of RNA from the mouse brain to the germline and offspring

**DOI:** 10.1101/686634

**Authors:** Elizabeth A. O’Brien, Kathleen S. Ensbey, Bryan W. Day, Paul A. Baldock, Guy Barry

## Abstract

**Background:** The traditional concept that heritability occurs exclusively from the transfer of germline-restricted genetics is being challenged by the increasing accumulation of evidence confirming the existence of experience-dependent transgenerational inheritance. However, questions remain unanswered as to how heritable information can be passed from somatic cells. Previous studies have implicated the critical involvement of RNA in heritable transgenerational effects and the high degree of mobility and genomic impact of RNAs in all organisms is an attractive model for the efficient transfer of genetic information.

**Results:** We hypothesized that RNA may be transported from a somatic tissue, in this case the brain, of an adult male mouse to the germline, and subsequently to offspring. To investigate this, we injected one hemisphere of the male mouse striatum with an AAV1/9 virus expressing human pre-MIR941 (MIR941). After 2, 8 and 16 weeks following injection, we used an LNA-based qPCR system to detect the presence of virus and human MIR941 in brain, peripheral tissues and offspring, from injected male mice mated with uninjected females. Virus was never detected outside of the brain. Verification of single bands of the correct size for MIR941 was performed using Sanger sequencing while quantitation demonstrated that a small percentage (∼ 1-8%) of MIR941 is transported to the germline and to offspring in about a third of the cases.

**Conclusions:** We show that somatic RNA can be transported to the germline and passed on to offspring, thereby providing additional evidence of a role for RNA in somatic cell-derived transgenerational effects.

## Introduction

Robust transgenerational inheritance has been observed in a diverse range of organisms, from protozoans and plants through to mammals [1–10]. These findings imply that a mechanism exists whereby experience-dependent information is transferred between somatic and germline tissues. Furthermore, multiple studies point to the potential existence of an RNA-mediated mechanism as a system for transferring genetic information [11–13] and potentially its stability across multiple generations [14]. The mode of transportation of RNAs may be via membrane-bound vesicles that are expelled from cells and contain an abundance of small RNAs [15–17], including miRNAs [18]. Long non-coding RNAs have also been indirectly postulated to be transferred to offspring due to experience-dependent alterations in the sperm [19–22]. We, therefore, postulated that somatically expressed RNAs may be transported to the germline allowing real-time heritability of somatic, experience-dependent alterations.

## Results

In order to test this hypothesis and trace genetic information from a somatic tissue to the germline in male mice, and to circumvent the issue of pervasive gene expression in sperm [23], we introduced a 72bp RNA encompassing a pre-isoform of a human-specific microRNA (MIR) 941, pre-MIR941-1 [24], via an adeno-associated virus (AAV) gene delivery system (Supplementary Fig. 1), into one hemisphere of the mouse striatum using stereotactic injection. The AAV serotype 1/9 (AAV1/9) vector, a single-stranded DNA packaging virus, has been shown to infect both neurons and astrocytes [25, 26]. The AAV1 vector was designed to also produce a single stranded DNA fragment of the rabbit β-globin gene which is used commercially for determining viral titre. We used this gene fragment to determine presence of the virus. Despite the fact that the AAV1 virus can be retrogradely transported via axons to the contralateral region [27] and the cerebellum [28], this system allowed us to restrict viral infection to the target site, and to discrete regions of the mouse brain while then specifically monitoring potential transport of virally-expressed pre-MIR941-1 outside of the brain using gene-specific PCR.

We initially sought to determine the specificity, reproducibility and detection limit using a locked nucleic acid (LNA) quantitative PCR (qPCR) system. Our methodological approach required a minimum length of the desired PCR product, hence the reason we targeted a 72bp pre-isoform transcript of MIR941. Henceforth, we will use the term ‘MIR941’ to describe the pre-MIR941-1 transcript. Using serial dilutions of positive control injection sites, qPCR reactions produced single bands of both MIR941 and rabbit β-globin as detected on an agarose gel (Supplementary Fig. 2). We found that we could detect MIR941 down to a 1:100 dilution (starting with 60ng RNA; Supplementary Fig. 2A) while the AAV1 virus could be detected down to a ∼ 1:15,000 dilution (starting with 60ng DNA; Supplementary Fig. 2B). We could not detect either MIR941 or rabbit β-globin fragment in control or mock-treated animals (Supplementary Fig. 3).

At two weeks post injection we found detectable MIR941 expression in the injection sites, however, no expression was detected in the contralateral site (Fig. 1; Supplementary Fig. 4). ‘Positive’ bands on a gel for all figures were supported by melt curve analysis and melting temperatures (see Supplementary Information). No expression was detected in the cerebellum or liver but, crucially, expression was detected in the lymph node and vas deferens and epididymis (V/E; Fig. 1; Supplementary Figs. 4, 5; Supplemental Information).

**Figure 1.**
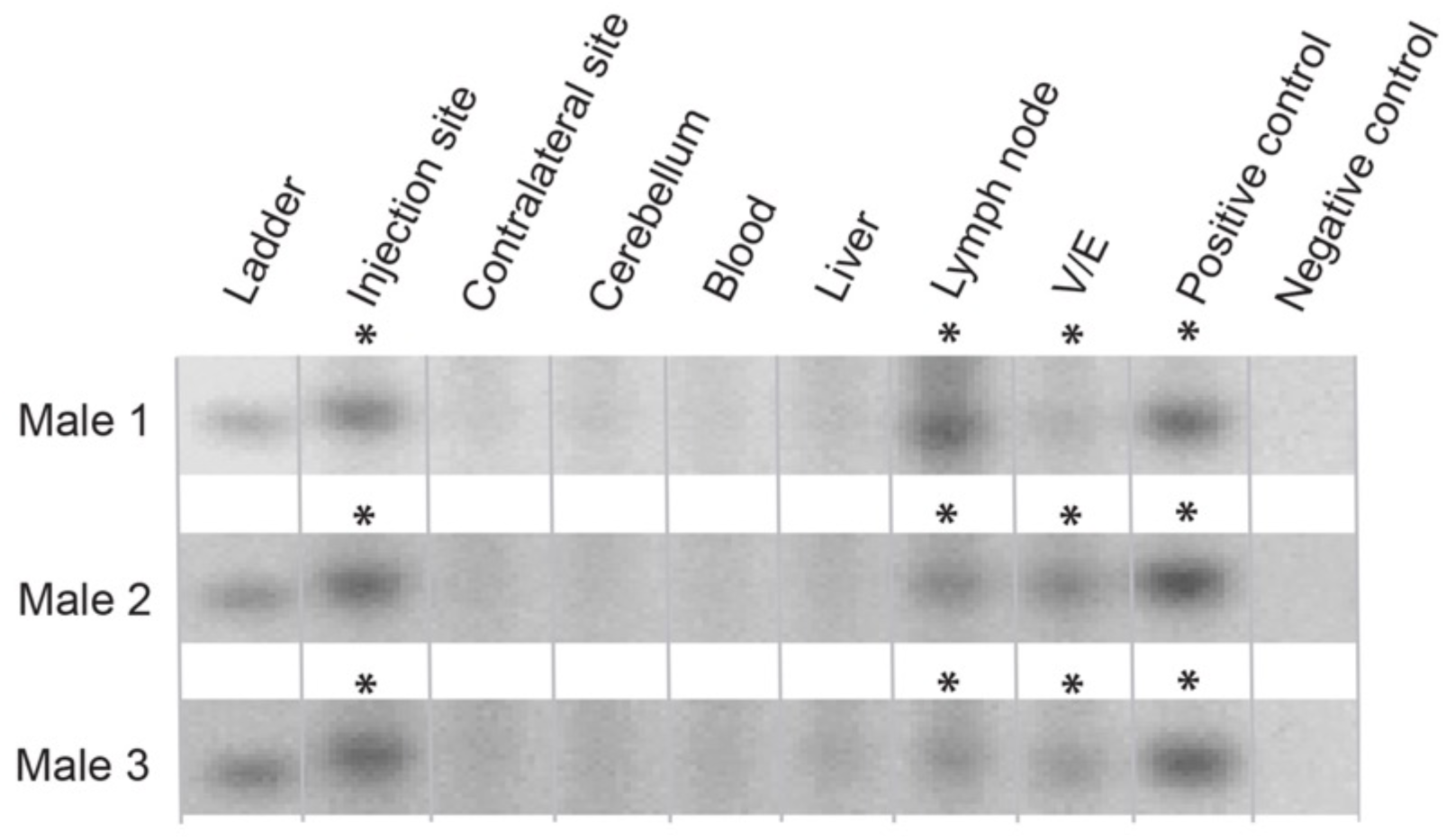
Detection of human MIR941 following 2 weeks post AAV1 injection into the striatum of male mice. An AAV1 viral vector, encoding human pre-MIR941-1 (MIR941), was injected into a single striatal hemisphere of three male mice. Following sacrifice after 2 weeks post-injection, MIR941 was detected, using gene-specific locked nucleic acid (LNA) quantitative PCR (qPCR), in the injection site, lymph node and vas deferens/epididymis (V/E). Human embryonic kidney (HEK) 293 cells were used as a positive control for MIR941 while a no template control served as a negative control. *Samples that were positive on the gel (single band of expected size) and were supported by melt curve analysis (Supplemental Information).

The evidence of MIR941 in the V/E, that contains mostly mature sperm, of the injected mice after two weeks prompted us to investigate whether this RNA may be transmitted to offspring. As a full cycle of spermatogenesis takes ∼ 5 weeks in mice, we mated male mice, after 6-8 weeks post-injection, with wildtype female mice. This would serve to increase the probability that brain-derived MIR941 had sufficient time to enter the testis and complete spermatogenesis. Male mice were sacrificed after 8 weeks and embryos were collected from wildtype female mice at 7.5 days post coitum (d.p.c.). Here we again detected MIR941 in samples from the injection and contralateral sites, cerebellum, lymph node and V/E (Fig. 2; Supplementary Figs. 6, 7). Crucially, we additionally detected MIR941 in 9 of the 18 embryos screened that originated from males treated with AAV1/MIR941. Viral presence was restricted to brain samples (Fig. 2; Supplementary Fig. 6).

**Figure 2.**
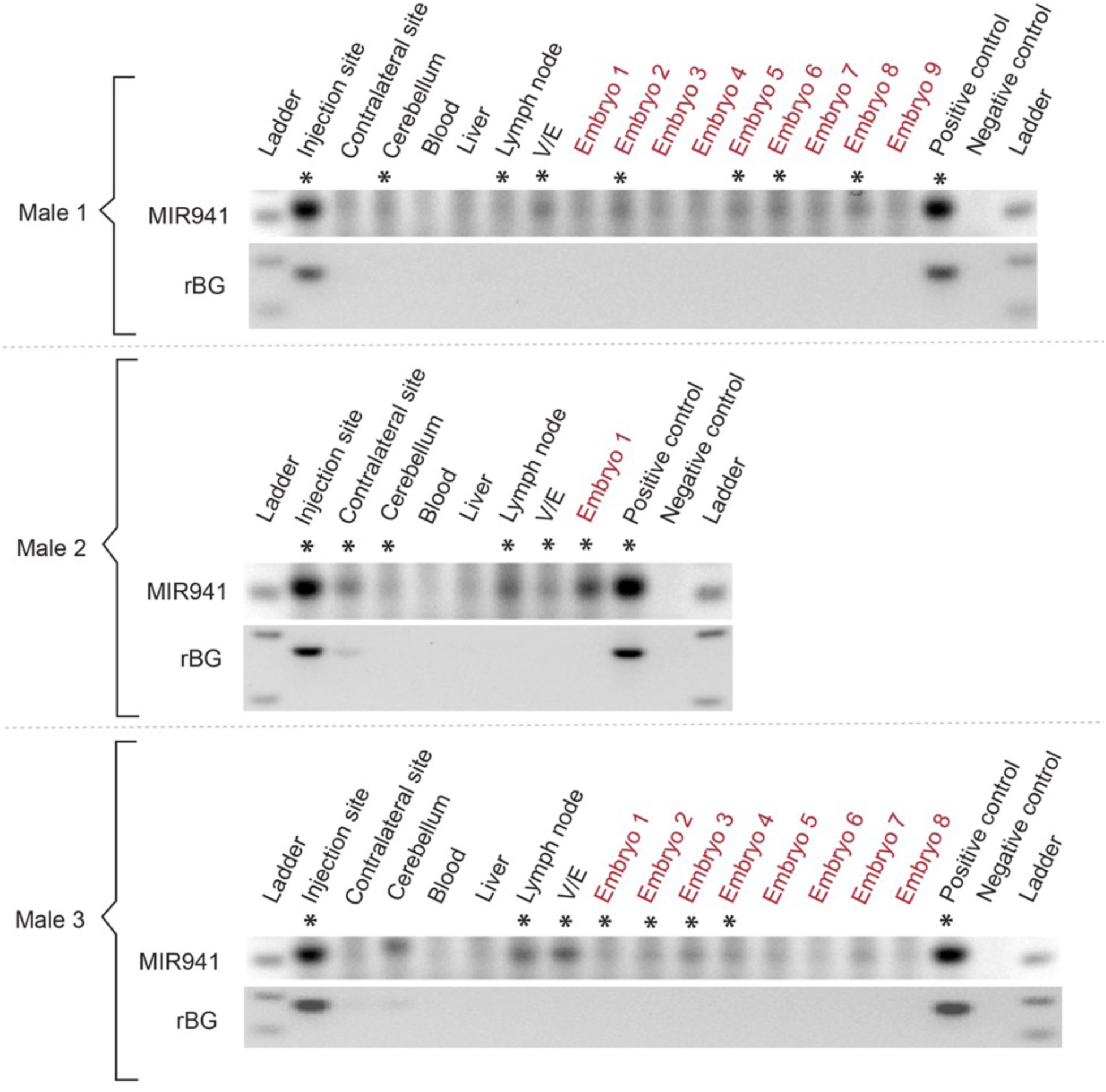
Detection of human pre-MIR941-1 in mouse embryos. AAV1-encoded pre-MIR941-1 (MIR941) was injected into one striatal hemisphere of three male mice. After 8 weeks post-injection, MIR941 was detected in injection and contralateral sites, cerebellum, lymph node and vas deferens/epididymis (V/E). Injected male mice were mated between 6-8 weeks following injection with wildtype female mice. Evidence of MIR941 was detected in 9 of the 18 embryos (7.5 d.p.c.) screened. Rabbit β-globin (rBG) fragment was only found in injection sites and one contralateral site. Positive control (HEK293), negative control (no template control). * Samples that were positive on the gel and were supported by melt curve analysis (Supplemental Information).

Although we could observe single bands of the correct size on the gels, we further validated all ‘positive’ bands using Sanger sequencing (Figs. 3A, B). Quantitative analysis, using UniSp6 as a spike-in control, showed that the amount of RNA found in the positive bands outside the brain were ∼ 1-2% of what is seen in the injection sites (Fig 3C).

**Figure 3.**
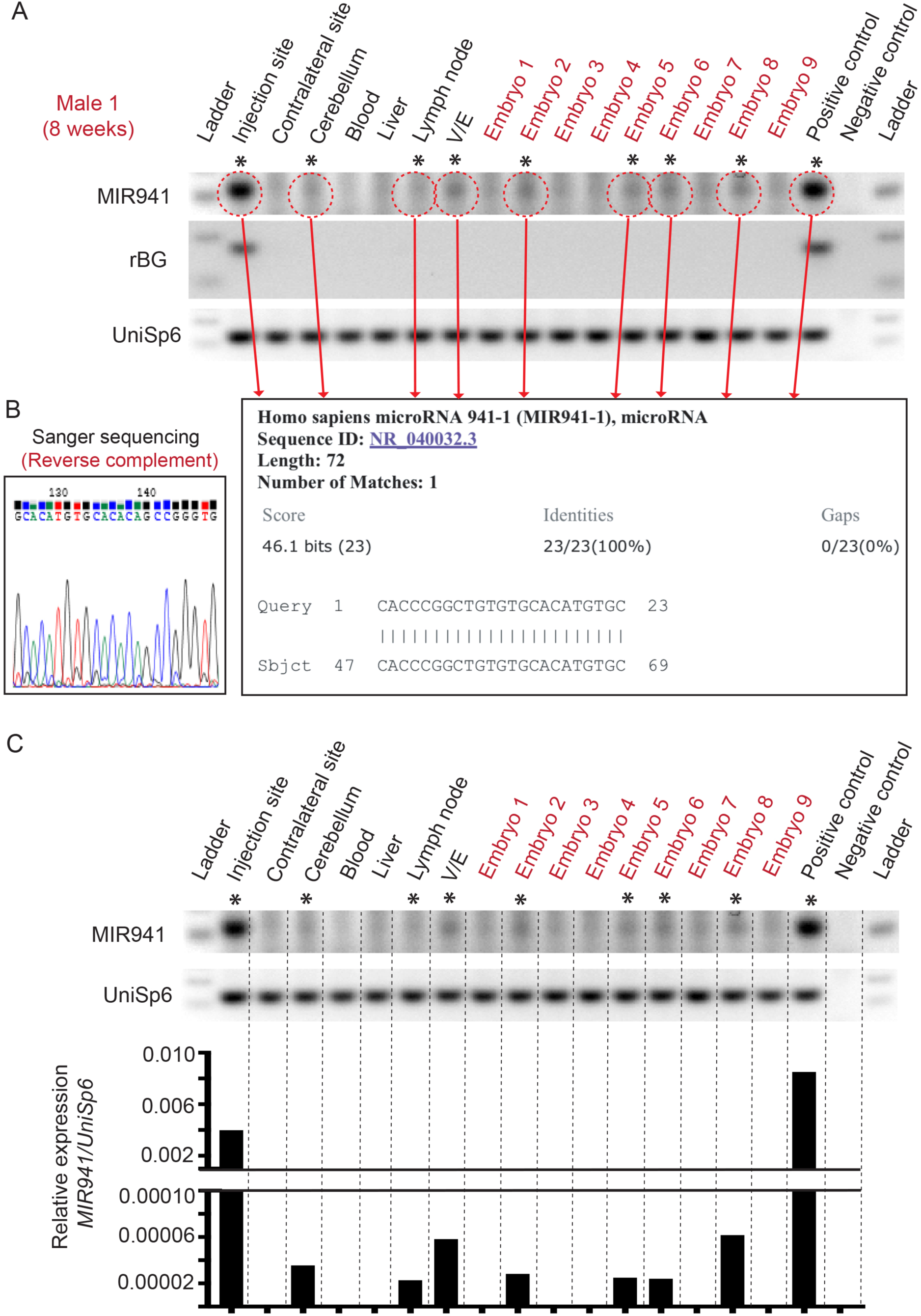
Sequencing and quantitation of ‘positive’ MIR941 LNA qPCR bands. (**A, B**) For example, we show that for Male 1 (Figure 2), all bands that were considered ‘positive’ for gel size and melt-curve analysis were also positive for human MIR941 following cloning and Sanger sequencing. (**C**) Quantitative analysis using UniSp6 spike-in controls demonstrate that the amount of MIR941 found to be positive outside of the brain is ∼ 1-2% of what is seen in the striatal injection sites.

We postulated that although we could detect MIR941 in some of the embryos, extending the time after injection and before mating may increase the detectability of MIR941 in embryos. Hence, we waited until 14 weeks post-injection to begin mating treated males with wildtype females and sacrificed the males at 16 weeks post-injection. Here we observed 17 out 60 embryos that were indeed positive for MIR941 (Fig. 4; Supplementary Fig. 8). These animals also reflected our previous results where MIR941 was detected in the injection and contralateral sites, cerebellum, lymph node and V/E but not in whole blood and liver. Viral presence was again only observed in discrete brain regions (Fig. 4; Supplementary Fig. 8).

**Figure 4.**
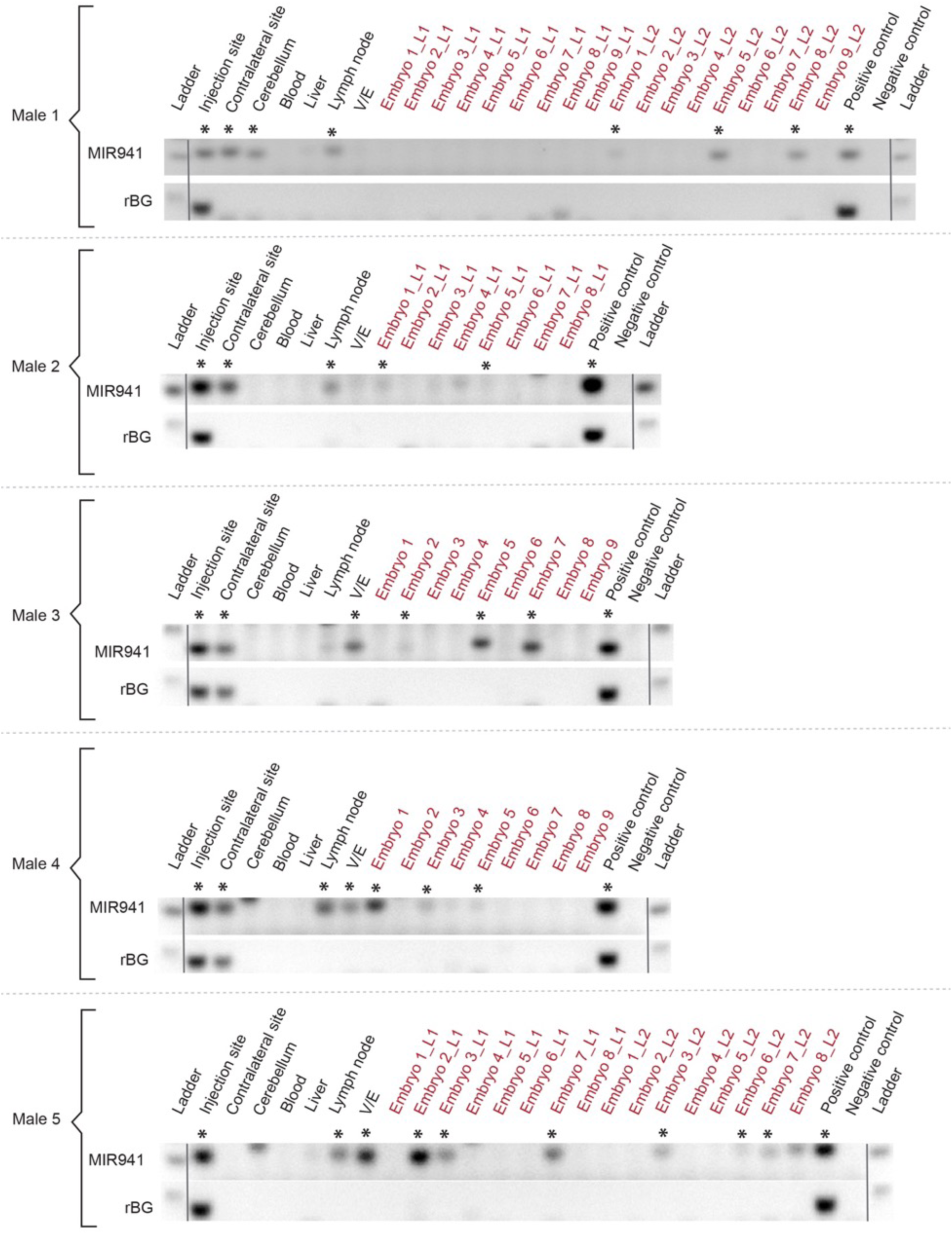
Detection of human pre-MIR941-1 in mouse embryos. AAV1-encoded pre-MIR941-1 (MIR941) was injected into one striatal hemisphere of five male mice. After 16 weeks post-injection, MIR941 was detected in injection and contralateral sites, cerebellum, lymph node and vas deferens/epididymis (V/E). Injected male mice were mated between 14-16 weeks post-injection with wildtype female mice and evidence of MIR941 was detected in 17 of the 60 embryos (7.5 d.p.c) screened. Rabbit β-globin (rBG) fragment was only found in injection and contralateral sites. Positive control (HEK293), negative control (no template control). * Samples that were positive on the gel and were supported by melt curve analysis.

Again, we further validated all ‘positive’ bands using Sanger sequencing (Figs. 5A, B). Quantitative analysis, using UniSp6 as a spike-in control, showed that the amount of RNA found in the positive bands outside the brain were ∼ 1-8% of what is seen in the injection sites (Fig 5C). These figures were higher than for 8 weeks and may reflect more time allowed for collection in peripheral tissues and germline.

**Figure 5.**
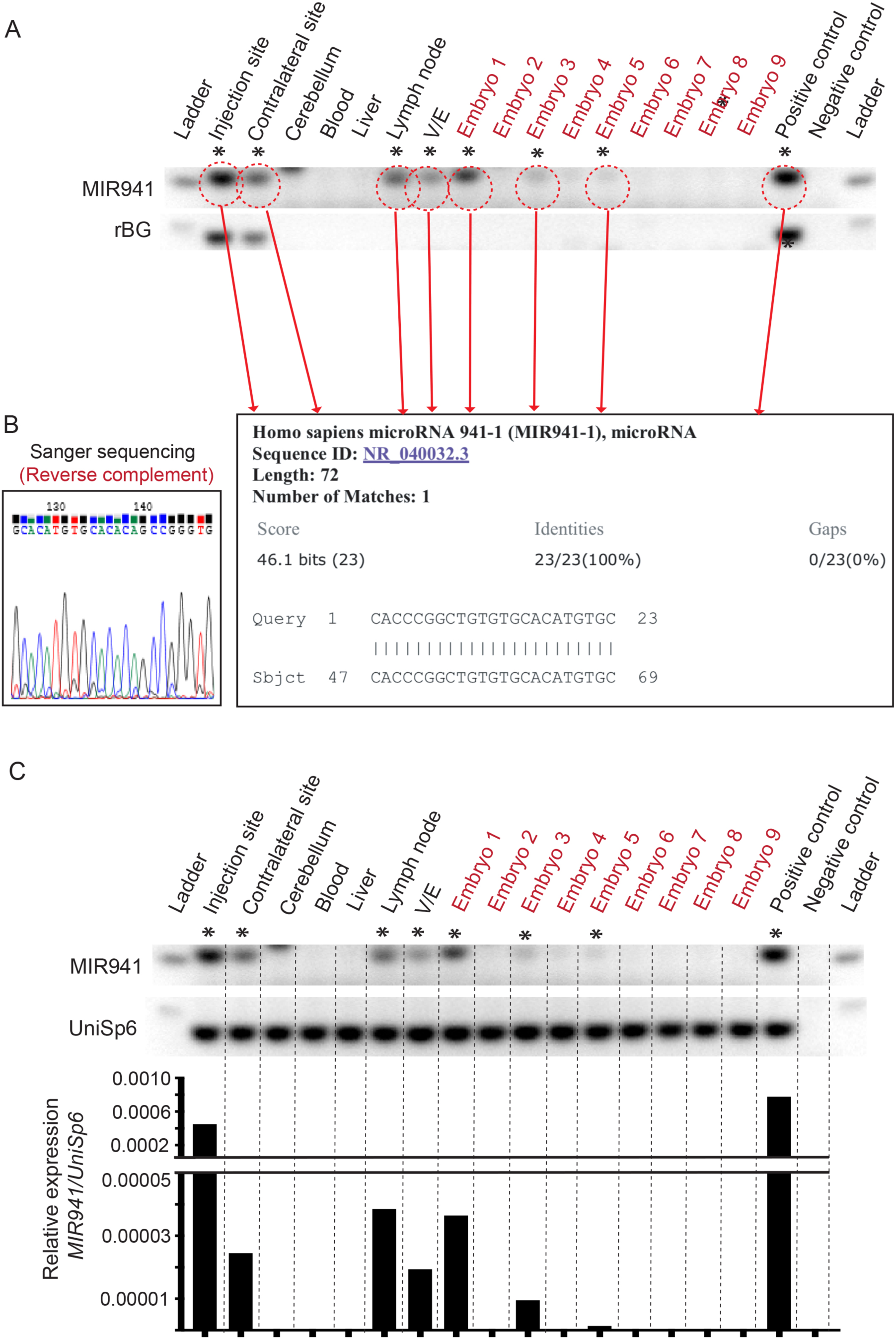
Sequencing and quantitation of ‘positive’ MIR941 LNA qPCR bands. (**A, B**) For example, we show that for Male 4 (Figure 4), all bands that were considered ‘positive’ for gel size and melt-curve analysis were also positive for human MIR941 following cloning and Sanger sequencing. (**C**) Quantitative analysis using UniSp6 spike-in controls demonstrate that the amount of MIR941 found to be positive outside of the brain is ∼ 1-8% of what is seen in the striatal injection sites.

The persistent presence of MIR941 in the lymph node does suggest that the mode of transport may be in both the lymphatic and circulatory systems considering their closely associated capillary transport of nutrients and small vesicles. We reasoned that MIR941 would be required to be contained in a vesicle so that degradation does not occur during transport, and vesicles are shared between the blood and the lymph due to their smaller size. However, whole blood, as opposed to the lymph node which collects lymph over a longer timeframe, may not be a suitable sample to detect what would be a very small amount of MIR941 at any one point, as blood flows faster than lymph. Therefore, in order to investigate whether MIR941 is present in the blood of the treated male mice we used a series of microcentrifuge and ultracentrifuge spins to isolate the fraction of whole blood that contains vesicles. We were not successful in detecting MIR941 in both the 2 or 8 week samples and only detected the presence of the miRNA in one of the 16 week samples in the ultracentrifuge fraction that is presumed to contain the highest concentration of vesicles (Supplementary Figs. 9A, B). No presence of the virus was found in any of these samples (Supplementary Fig. 9C).

## Discussion

Our data verify that RNAs are able to be physically transmitted from a somatic tissue, in this case the brain, to the germline and to offspring in male mice. The mode of transport for the RNA via the lymph and bloodstream is likely to be vesicles, which would provide a secure environment to avoid degradation and increase stability [29, 30]. Although we could only find one case where our detection system captured the RNA in the vesicle-containing fractions of ultracentrifuged whole blood (Supplementary Fig. 9), this may serve to only highlight current detection limitations. Therefore, we are unable to confirm the exact prevalence because we may not be capturing all positive samples, since our experimental design purposely imposed a low level of viral infection so as not to flood the endogenous system. However, combined data from both 8 and 16 week post-injected males and their respective offspring still showed a 33% heritability rate. Furthermore, our thorough verification and quantitation of positive qPCR results from multiple animals and timeframes, combined with ensuring that the virus does not escape the brain, leads us to confidently conclude that this process does indeed occur at least in the mouse. The demonstration that a third of embryos contain the MIR941 transcript after 8 and 16 weeks post-injection does suggest that RNA transfer reflects a meaningful mechanism for transgenerational epigenetic effects and sets the foundation for direct measurements of the extent of RNA-based somatic inheritance in different species.

## Conclusion

Our results provide further evidence for an essential role of mobile RNA in the transmission of somatically-derived epigenetic transgenerational effects.

## Materials and Methods

### AAV vector

The plasmid preparation and viral vector production was performed at Penn Vector Core, University of Pennsylvania. The plasmid vector, pENN.AAV.CB7.CI.MIR941.rBG, was generated by inserting pre-MIR941-1 (Sequence: TGTGGACATGTGCCCAGGGCCCGGGA CAGCGCCACGGAAGAGGACGCACCCGGCTGTGTGCACATGTGCCCA) into the cloning plasmid pAAV.CB7.rBG. Briefly, the CB7 promoter - composed of CMV enhancer, beta-actin promoter and a chimeric intron- drives the expression of the pre-MIR941-1 transcript. The viral vector derived from the plasmid, AAV1.CB7.CI.MIR941.rBG, was produced, titred and supplied in PBS containing 0.001% Pluronic F68.

### Animals

All animal procedures were approved by the QIMRB Animal Ethics committee, ethics #A1612-622M. C57BL/6J (Jax code 000664) were obtained from Animal Resource Centre (Canning Vale).

### Brain injections

Male C57BL/6J aged 6-7 weeks were administered analgesia (Meloxicam 5mg/kg) 30 minutes prior to surgery. Animals were anaesthetized using inhaled isoflurane (2%) and once anaesthetized were transferred onto the stereotactic device with nose cone attachment for continuous isoflurane administration. Once the mouse was adequately anaesthetized by evidence of pedal withdrawal reflex, the animal’s head was clipped, disinfected using an alcohol swab, the skin was incised and a 2mm drill hole was made and a solution (1-2ul) containing AAV vector was injected (Hamilton syringe) into the lateral portion of the striatum. Injections were made over a period of 3 minutes to ensure optimal parenchymal compliance. The micro-injector needle was retracted, the bur hole was plugged with bone wax, a ‘splash block’ of Bupivacaine (5mg/kg) was applied to the skull/periosteum and the skin closed with tissue adhesive (VetBond). Animals were returned to their cages, given free access to food and water and monitored daily for any signs of neurological dysfunction. On the day following surgery the animals were administered another dose of analgesia (Meloxicam 5mg/kg).

### Timed matings

Wildtype C57BL/6J females aged 6-8 weeks were set up overnight with a male. The following morning presence of a vaginal plug assumed that mating/fertilization occurred. We determine that embryos at this stage are aged 0.5 days post coitum (d.p.c).

### Collection of Embryos

Post implantation embryos at 7.5 d.p.c were collected from euthanized pregnant females. The uterine horn was dissected out and placed in ice cold PBS. The embryos were separated by cutting between implantation sites along the uterine horn. The surrounding muscle layer is peeled back to expose the decidua. Individual embryos still encased in decidua tissue were flash frozen on dry ice and stored at - 80°C until homogenization for nucleic acid extraction.

### Dissection for tissue collection

For tissue collection, animals were anaesthetized and then cardiac puncture was performed. Once blood was collected, cervical dislocation was performed. The following tissues were collected: injection site (striatum), contralateral striatum, cerebellum, inguinal lymph nodes, liver, vas deferens and epididymis (V/E). Dissected tissue was collected into Eppendorf tubes and flash frozen on dry ice. Storage at - 80°C prior to nucleic acid extraction.

### Blood and Plasma samples

Immediately after collection of blood, 0.1ml of blood was diluted in 0.15ml nuclease free water (Invitrogen #10977015) and 0.75ml TRIzol LS Reagent (ThermoFisher #10296010) was added. Samples were immediately flash frozen on dry ice and stored at -80°C prior to nucleic acid extraction. The remaining blood collected from cardiac puncture was placed in MiniCollect K2E K2EDTA 0.5ml (Greiner Bio-One #450480). Manufacturer’s instructions were followed to collect the resulting plasma. Plasma was stored at -80°C.

### Ultracentrifugation of Plasma Samples

Plasma samples were thawed on ice. Three successive centrifugation steps were performed. The pellet at each step was collected and resuspended in TRIzol Reagent (ThermoFisher #15596026), while the supernatant was centrifuged again. The first centrifugation step was performed at 3,000 x g for 10 minutes at 4°C. The second centrifugation step performed was 12,000 x g for 45 minutes at 4°C. The final centrifugation step performed was 110,000 x g for 70 minutes at 4°C in a Hitachi Micro Ultracentrifuge CS150NX. Samples were immediately flash frozen on dry ice and stored at -80°C prior to nucleic acid extraction.

### Homogenization of tissue samples

Tissue samples were transferred while still frozen into screw capped tubes containing approximately 8-10 1.4mm zirconium oxide beads (Bertin Technologies KT03961-1-103.BK). Samples had appropriate amounts of TRIzol Reagent (ThermoFisher #15596026) added as per manufacturer’s instructions. Samples were homogenized in Precellys 24 (ThermoFisher) twice at 6,000rpm for 30 seconds with a hold period in between. The homogenized sample was transferred into an Eppendorf tube to remove the beads. The samples were then centrifuged 12,000 x g for 5 minutes at 4°C. The supernatant was collected in a new Eppendorf tube. Samples were stored at -80°C prior to nucleic acid extraction.

### Extraction of nucleic acids (RNA and DNA)

RNA and DNA were extracted from each sample. All samples had appropriate amounts of chloroform added to the TRIzol Reagent/TRIzol LS Reagent as per manufacturer’s instructions. The aqueous layer that formed after centrifugation at 12,000 x g for 15 minutes at 4°C was collected and combined with 1.5 x volume 100% ethanol. Purification of microRNA and total RNA from this fraction was performed using miRNeasy Mini/Micro Kit (Qiagen #217004/#217084) as per manufacturer’s instructions. On-column DNase digestion was performed using the RNase-Free DNase Set (Qiagen #79254).

After the aqueous phase was collected for extraction of RNA, the remaining interphase and phenol chloroform phase was used to extract DNA with the addition of Back Extraction Buffer (BEB - 4M guanidine thiocyanate, 50mM sodium citrate, 1M Tris), 0.25ml BEB per 0.5ml TRIzol. The solution was shaken vigorously for 15 seconds, incubated at room temperature for 10 minutes and then centrifugated at 12,000 x g for 15 minutes at 4°C. The resulting aqueous layer was collected, mixed with 1 x volume of 100% ethanol, and the DNA was purified using DNeasy Blood & Tissue Kit (Qiagen #69506) as per manufacturer’s instructions.

RNA/DNA was quantified using NanoDrop Lite (ThermoFisher).

### cDNA synthesis

cDNA synthesis using the miRCURY LNA miRNA PCR Starter Kit (Qiagen #3390320) was performed as per manufacturer’s instructions. One first-strand cDNA synthesis reaction per sample was performed. 200ng total RNA per 10µl cDNA synthesis reaction was performed for all samples except for plasma samples where 10ng total RNA per 10µl cDNA synthesis reaction was used. Each reaction included UniSp6 spike-in control. Protocol as per manufacturer’s instructions was followed.

### Real-time PCR amplification with LNA-enhanced primers

Real-time PCR was performed using components from miRCURY LNA miRNA PCR Starter Kit. All runs were performed using the QuantStudio5 Real-Time PCR System (Applied Biosystems) in “fast” mode using ROX as the passive reference dye. Each reaction had a total volume of 10µl.

### hsa-MIR-941miRCURY LNA miRNA PCR Assay

This was a pre-designed, validated assay to target hsa-MIR-941 (Qiagen product# 339306, catalogue# YP00204574). Each 10µl reaction was made up of the following: 5µl 2 x miRCURY SYBR Green Master Mix, 0.05µl ROX Reference Dye, 1.95µl PCR primer mix, 3µl of cDNA template (diluted 1:10). The PCR was initially heat activated at 95°C for 2 minutes, followed by 40 cycles of denaturation (95°C for 10 seconds) and combined annealing/extension (60°C for 60 seconds).

### UniSp6 miRCURY LNA miRNA PCR Assay

This was a pre-designed, validated assay to target UniSp6 (Qiagen product# 339306, catalogue# YP00203954) which is a cDNA synthesis/PCR control. Each 10µl reaction was made up of the following: 5µl 2 x miRCURY SYBR Green Master Mix, 0.05µl ROX Reference Dye, 1µl PCR primer mix, 0.95µl nuclease-free water, 3µl of cDNA template (diluted 1:60). The reaction was initially heat activated at 95°C for 2 minutes, followed by 40 cycles of denaturation (95°C for 10 seconds) and combined annealing/extension (60°C for 60 seconds). All reactions were positive for UniSp6.

### Rabbit β-Globin qPCR

Custom qPCR LNA primers were designed through Exiqon prior to Qiagen’s acquisition of the company. They were designed to target rabbit β-globin poly A tail in the virus. Each 10µl reaction was made up of the following: 5µl 2 x miRCURY SYBR Green Master Mix, 0.05µl ROX Reference Dye, 1µl PCR primer mix, 0.95µl nuclease-free water, 3µl of DNA template (1ng/λ). The reaction was initially heat activated at 95°C for 2 minutes, followed by 35 cycles of denaturation (95°C for 10 seconds) and combined annealing/extension (60°C for 60 seconds). Primers sequences: Forward – TATGGGGACATCATGAAGC; Reverse - CCAACACACTATTGCAATGA

### Gels

qPCR products were run on 3% TAE agarose gels containing 1 x Biotium GelRed NA Stain (Fisher Biotech #41003) at 100V for 90 mins prior to imaging/image capture.

### Cloning and sequencing of qPCR product

qPCR for hsa-miR-941 was performed as per the conditions outlined above. Samples were determined to be positive based on Ct value and melt curve analysis. These positive qPCR products were purified using a MinElute PCR Purification Kit (Qiagen #28004) as per manufacturer’s instructions. A 20ul ligation reaction was set up which contained 9ul of purified insert, 1ul (25ng) pCR2.1 linearized vector (ThermoFisher #K202020), 2ul of 10 x T4 DNA Ligase Buffer (New England Biolabs # M0202S), 1ul (400Units) of T4 DNA Ligase (New England Biolabs # M0202S), 7ul nuclease free water. The reaction was conducted at 16°C for 16 hours. Transformation reactions were set up using 5ul of ligation reaction and 50ul chemically competent *E.coli* cells (New England Biolabs # C3040I) and performed as per manufacturer’s instructions. 40ul of 20mg/mL X-gal (5-bromo-4-chloro-3-indolyl-β-D-galacto-pyranoside – Sigma Aldrich B44252) and 40ul of 100mM IPTG (isopropyl β-D-1-thiogalactopyranoside – AppliChem A4773) was mixed 1:1 prior to plating on the surface LB Agar plates containing 100ug/mL Ampicillin (Sigma Aldrich #A9518) and this was allowed to dry prior to the transformation reaction being plated. The plates were incubated 30°C overnight and the following day white colonies were selected for further analysis. Each white colony was picked and inoculated into 3mL LB broth containing 100ug/mL Ampicillin (Sigma Aldrich #A9518). These liquid cultures were grown at 30°C with shaking (250 rpm) for 20 hours. The cultures then had the DNA extracted from them using the QIAprep Spin Miniprep Kit (Qiagen #27104) and the resulting DNA was quantified using NanoDrop Lite (ThermoFisher). For sequencing using the BigDye Terminator v3.1 Cycle Sequencing Kit (ThermoFisher), 150ng of vector and 3pmol of primer (M13F) were used per reaction as per manufacturer’s instructions. The reaction was cleaned up and run on the 3130xl Genetic Analzyer (ThermoFisher). The resulting sequence was used as input for BLAST to confirm insert identity.

## Declarations

The authors declare no competing interests

## Acknowledgments

We thank D. McNeilly for his technical expertise and animal handling.

## Author Contributions

G.B. and P.A.B. conceived the study. G.B. acquired funding and designed the research. K.S.E. performed the stereotactic injections and dissections. E.A.B. performed dissections, all DNA/RNA extractions and all molecular analysis. G.B., P.A.B. and B.W.D. contributed resources. E.A.B. and G.B. wrote the manuscript, all other authors contributing to revision of the manuscript. G.B. supervised and provided oversight for the project.

## Supplementary Information

**Melting temperatures (T**_**m**_**) for (*) positive calls for MIR941 on the gels (Figures 1-3).** The following represent positive calls made in the main text as an asterix (*) in figures 1-3. For a ‘positive’ call, there needed to be a correct size band on the gel (products generated by LNA qPCR), a graphical ‘positive’ via melt curve analysis and T_m_ values that were consistent with positive control values in each run.

*(All other T*_*m*_ *values were either significantly divergent from these ‘positive’ values or ‘undetermined’)*

### Figure 1: 2 weeks

T_m_ (Injection sites): (Male 1) 72.953, (Male 2) 72.834, (Male 3) 72.834

T_m_ (Lymph nodes): (Male 1) 72.114, (Male 2) 72.114, (Male 3) 72.354

T_m_ (Vas/Epididymis): (Male 1) 72.354, (Male 2) 72.474, (Male 3) 71.994

T_m_ (Positive controls): (Male 1) 73.073, (Male 2) 73.073, (Male 3) 72.953

### Figure 2: 8 weeks

T_m_ (Injection sites): (Male 1) 72.867, (Male 2) 72.986, (Male 3) 72.629

T_m_ (Contralateral site): (Male 2) 72.509

T_m_ (Cerebellum): (Male 1) 72.271, (Male 2) 73.462

T_m_ (Lymph nodes): (Male 1) 72.867, (Male 2) 72.033, (Male 3) 71.914

T_m_ (Vas/Epididymis): (Male 1) 72.986, (Male 2) 71.795, (Male 3) 72.509

T_m_ (Positive controls): (Male 1) 73.224, (Male 2) 73.105, (Male 3) 73.105 T_m_ (Embryos):

- Male 1 - (Embryo 2) 73.224, (Embryo 5) 73.224, (Embryo 6) 72.748, (Embryo 8) 72.629
- Male 2 - (Embryo 1) 72.629
- Male 3 - (Embryo 1) 72.309, (Embryo 2) 72.309, (Embryo 3) 73.343, (Embryo 4) 72.748

### Figure 3: 16 weeks

T_m_ (Injection sites): (Male 1) 71.636, (Male 2) 71.516, (Male 3) 71.796, (Male 4) 71.396, (Male 5) 71.396.

T_m_ (Contralateral site): (Male 1) 71.996, (Male 2) 72.236, (Male 4) 71.516

T_m_ (Cerebellum): (Male 1) 70.197

T_m_ (Lymph nodes): (Male 1) 71.156, (Male 2) 70.916, (Male 4) 73.548, (Male 5) 71.156

T_m_ (Vas/Epididymis): (Male 3) 73.548, (Male 4) 71.156, (Male 5) 71.156

T_m_ (Positive controls): (Male 1) 71.636, (Male 2) 71.636, (Male 3) 71.796, (Male 4) 71.636, (Male 5) 71.636

T_m_ (Embryos):

- Male 1 - (L2 Embryo 1) 71.156, (L2 Embryo 5) 71.636, (L2 Embryo 8) 71.756
- Male 2 - (Embryo 1) 71.036, (Embryo 5) 71.156
- Male 3 - (Embryo 2) 71.445, (Embryo 5) 72.964, (Embryo 7) 72.146
- Male 4 - (Embryo 1) 71.996, (Embryo 3) 72.356, (Embryo 5) 71.276
- Male 5 - (L1 Embryo 2) 71.516, (L1 Embryo 3) 70.916, (L1 Embryo 7) 70.916, (L2 Embryo 3) 71.036, (L2 Embryo 6) 71.756, (L2 Embryo 7) 71.036

## Supplementary Figures

**Supplementary Figure 1.**
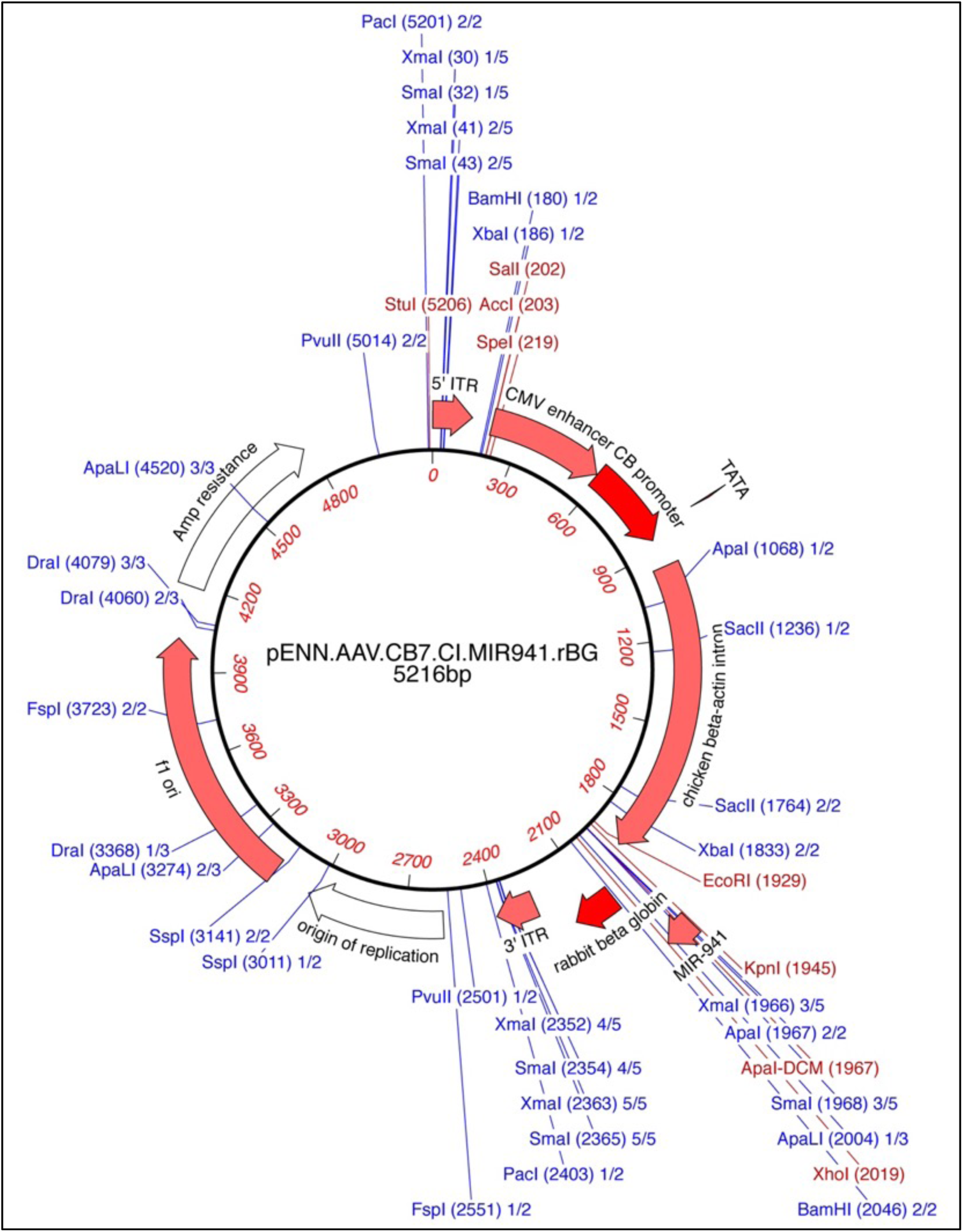
pENN.AAV.CB7.CI.MIR941.rBG. Plasmid produced for viral vector production by Penn Vector Core.

**Supplementary Figure 2.**
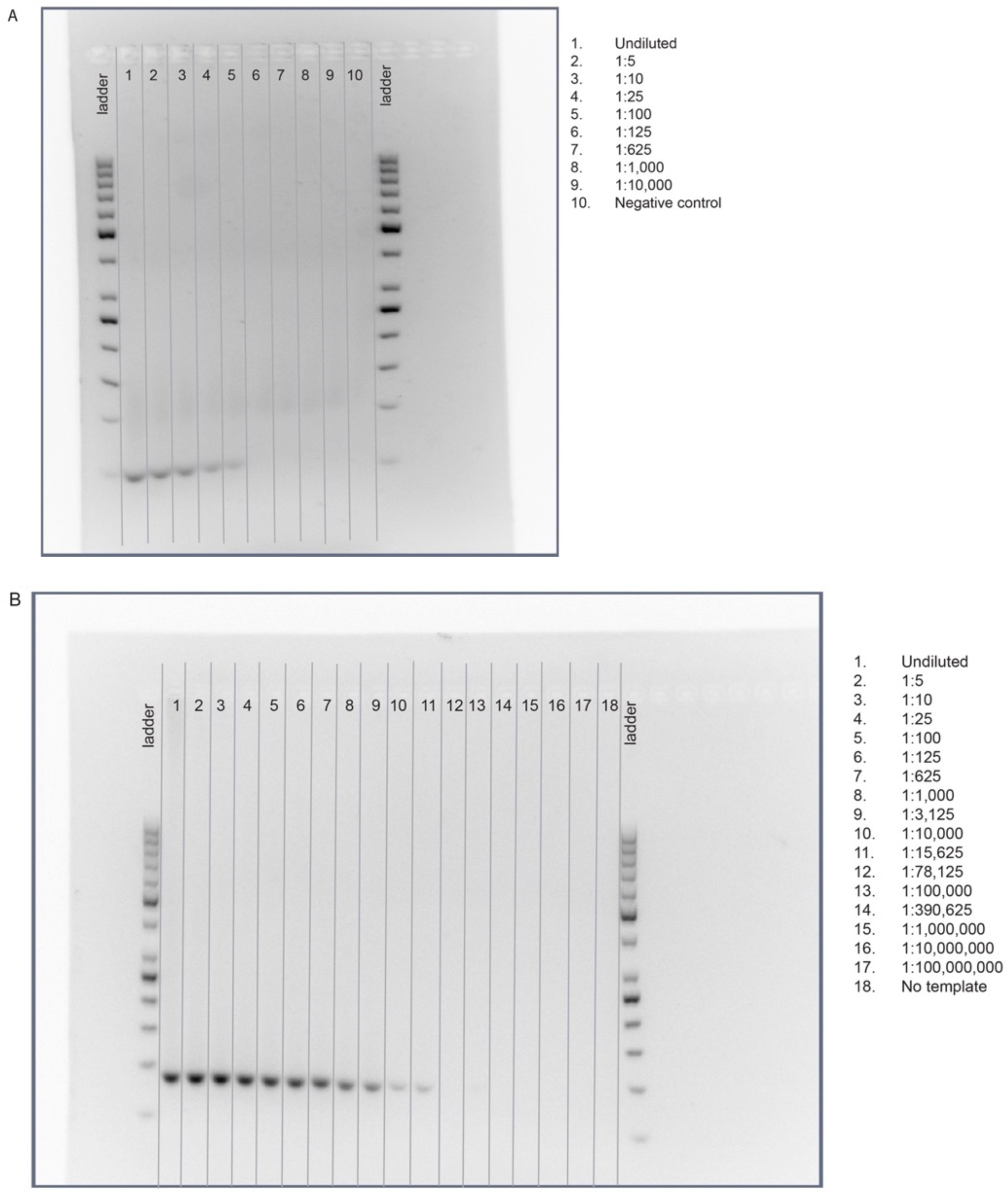
Detection of (**A**) MIR941 and (**B**) a rabbit β-globin fragment in treated adult male mice. Positive control striatal injection sites from adult male mice were used to detect single bands using a gene-specific locked nucleic acid (LNA) quantitative PCR (qPCR) system.

**Supplementary Figure 3.**
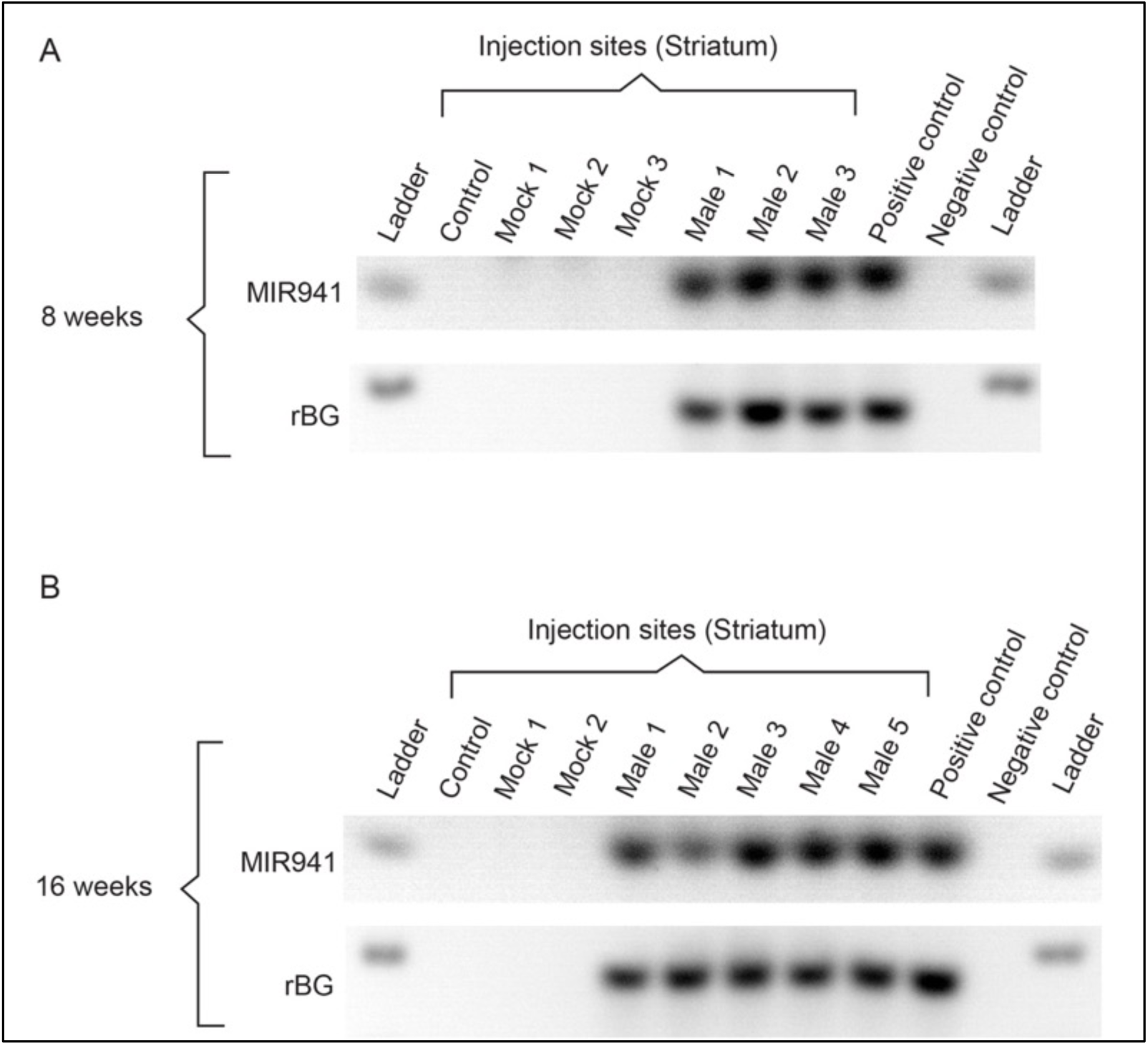
No detection of MIR941 or rabbit β-globin fragment in control (uninjected) or mock-treated (saline-injected) animals at either (**A**) 8 weeks or (**B**) 16 weeks post-injection.

**Supplementary Figure 4.**
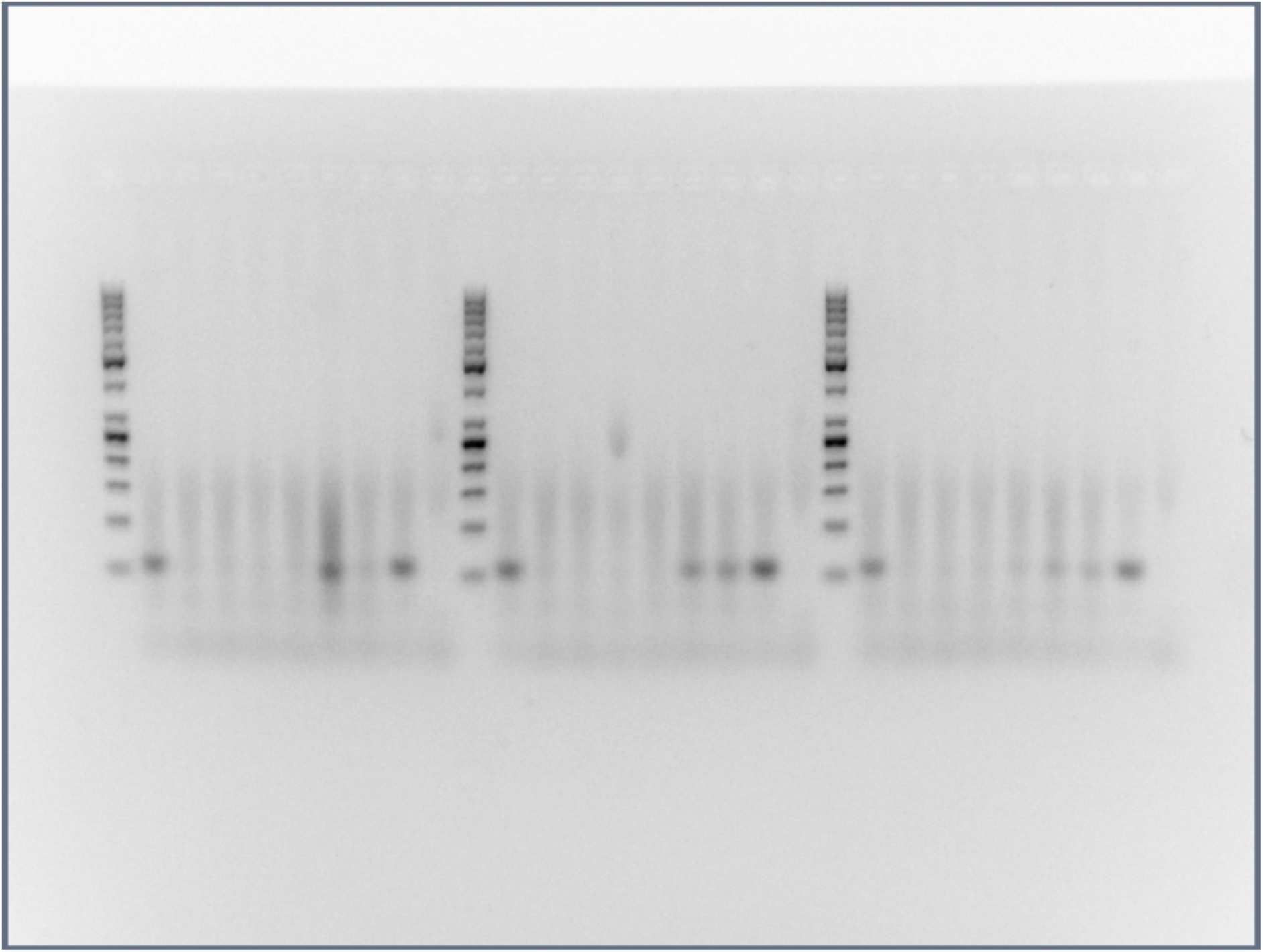
Full size, unedited gels used for Figure 1 in the main text.

**Supplementary Figure 5.**
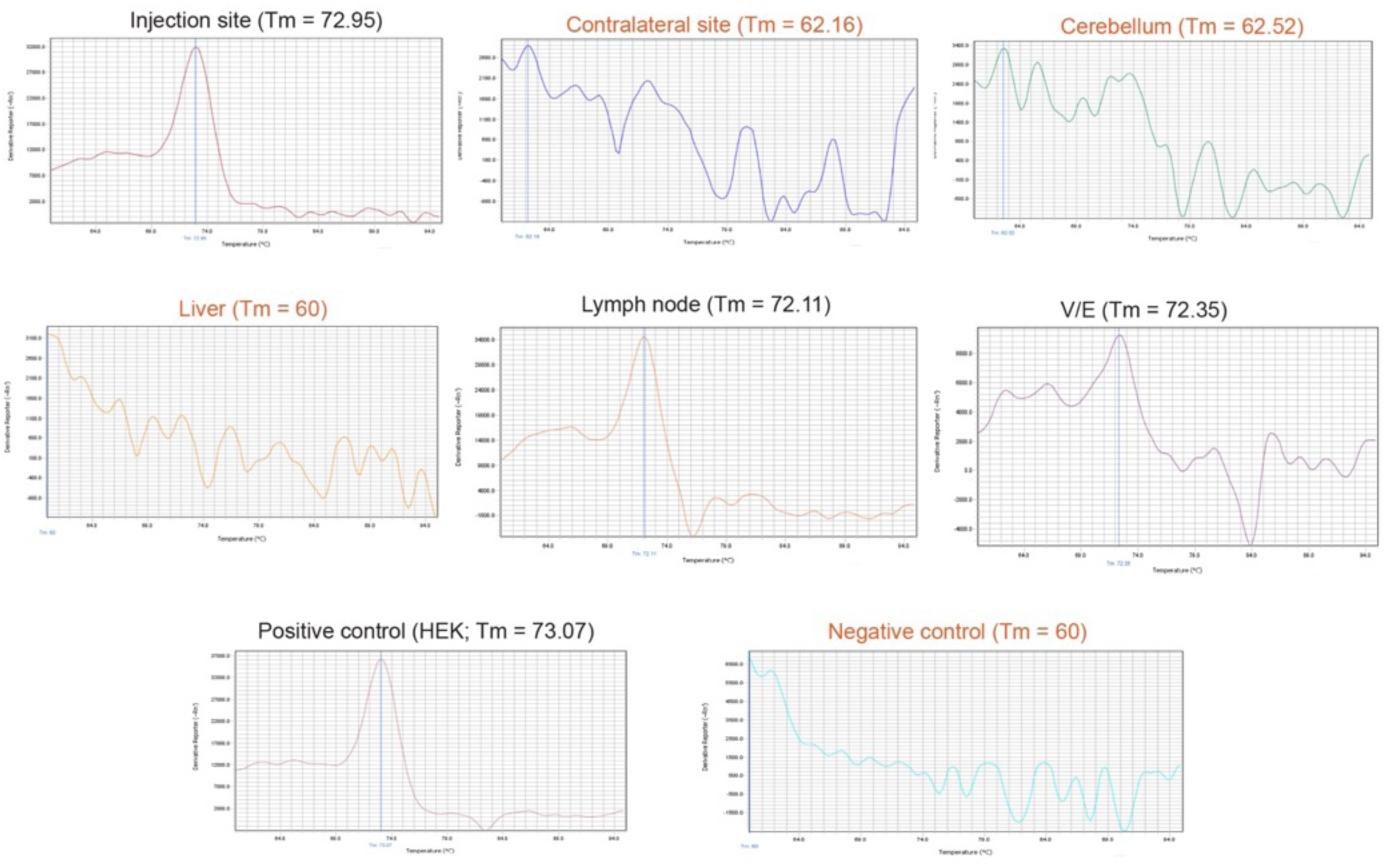
Representative full panel of melt curves for Week 2, Male 1 showing MIR941 LNA qPCR. Positive (black writing) and negative (red writing) melt curves and associated melting temperatures (T_m_) are shown and correspond to the bands in Figure 1. We have used an asterix (*) to denote a positive band of correct size on the gel that also had positive melt curves and T_m_ values as shown in this figure.

**Supplementary Figure 6.**
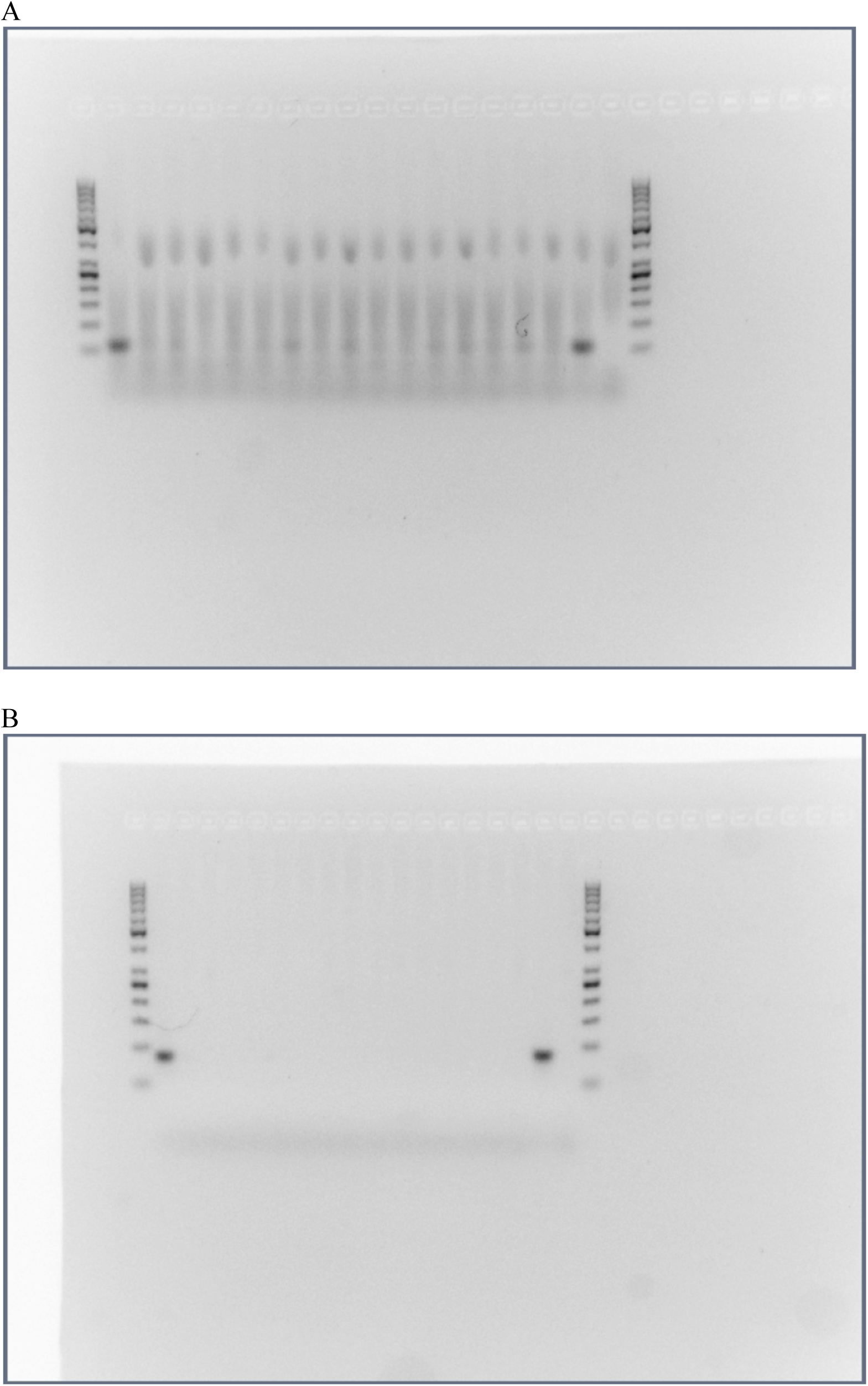
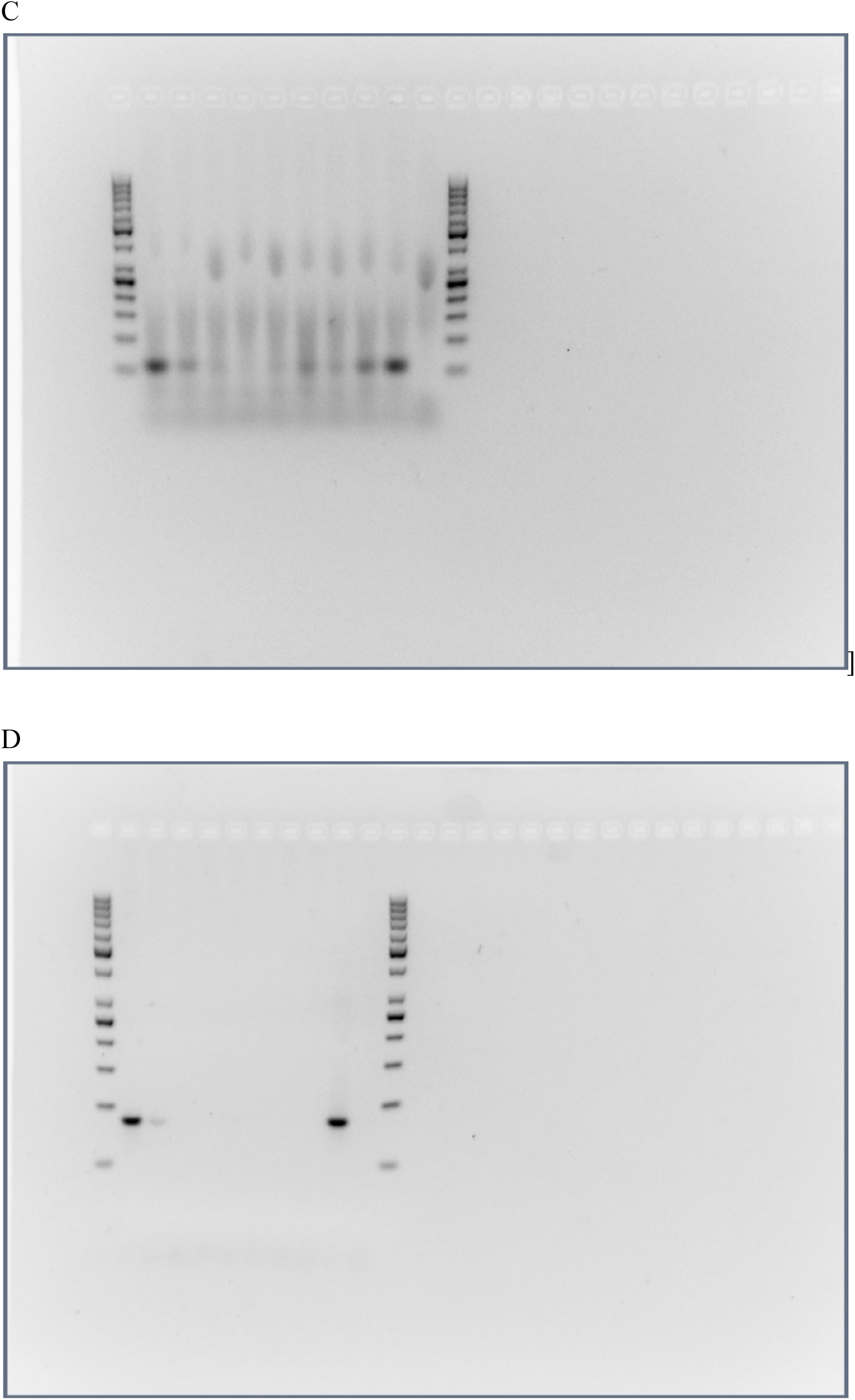
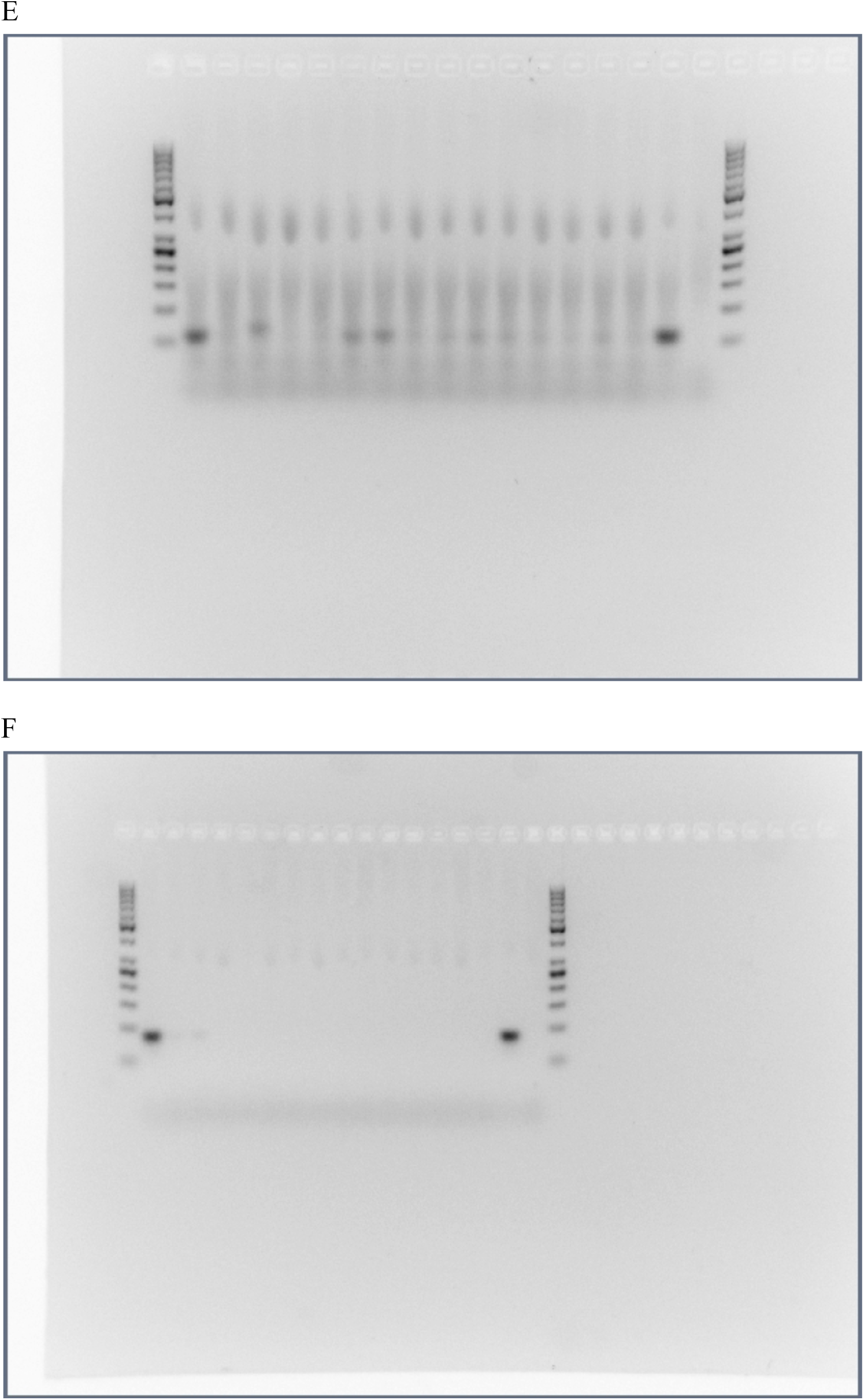
Full size, unedited gels used for Figure 2 in the main text. (**A**) MIR941-Male 1, (**B**) Rabbit β-globin fragment - Male 1, (**C**) MIR941-Male 2, (**D**) Rabbit β-globin fragment - Male 2, (**E**) MIR941-Male 3, (**F**) Rabbit β-globin fragment - Male 3.

**Supplementary Figure 7.**
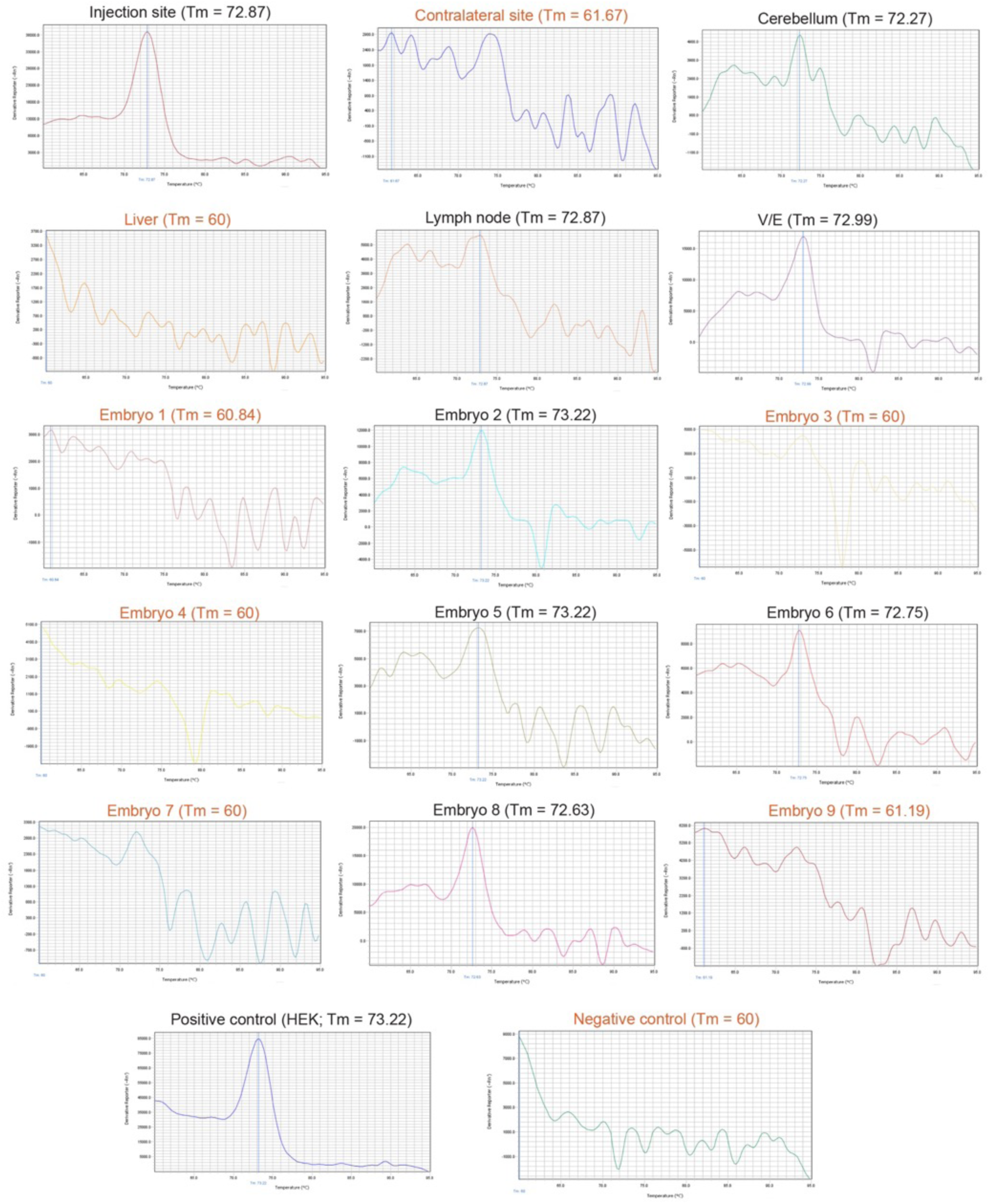
Representative full panel of melt curves for Week 8, Male 1 showing MIR941 LNA qPCR. Positive (black writing) and negative (red writing) melt curves and associated melting temperatures (T_m_) are shown and correspond to the bands in Figure 2. We have used an asterix (*) to denote a positive band of correct size on the gel that also had positive melt curves and T_m_ values as shown in this figure.

**Supplementary Figure 8.**
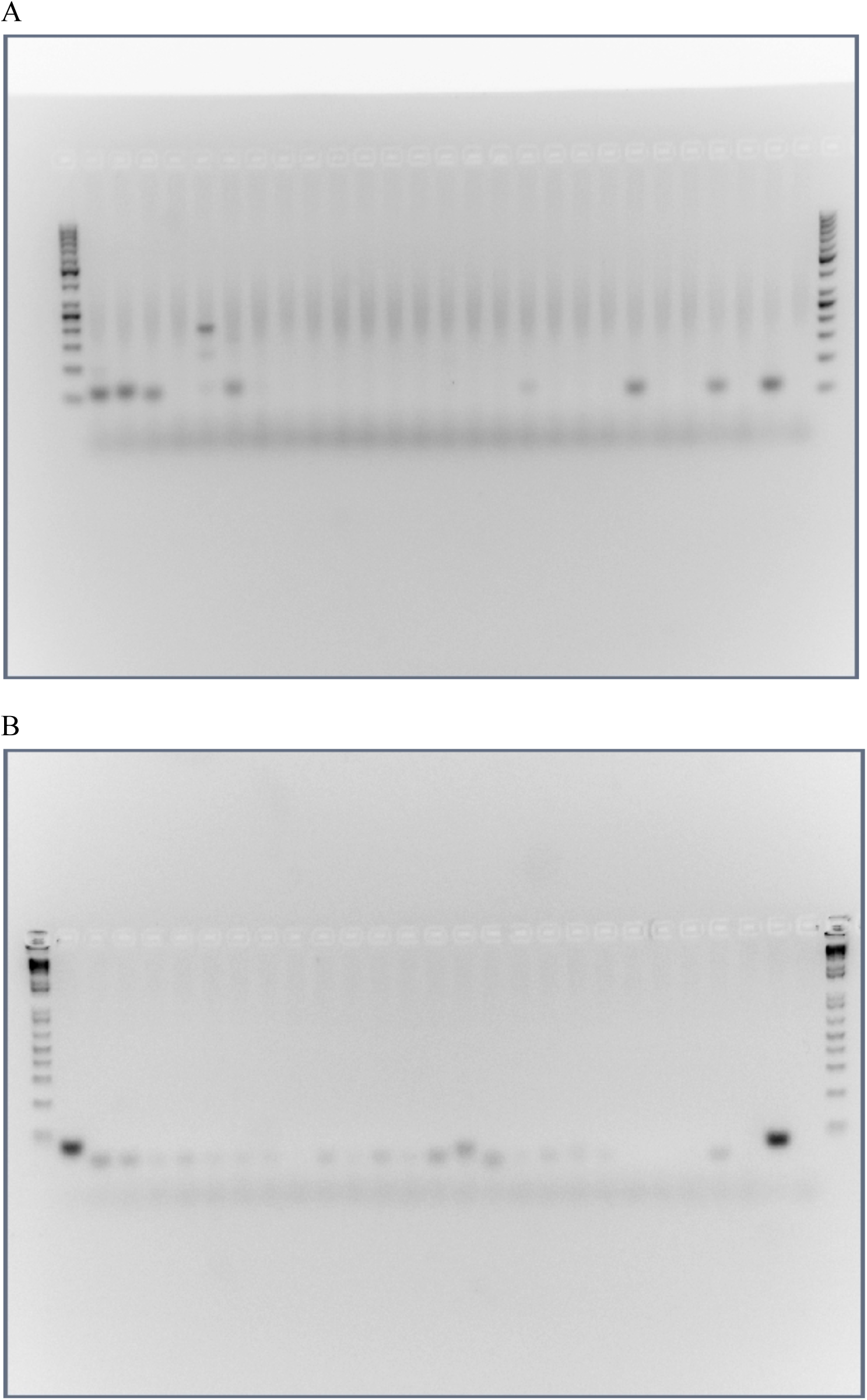
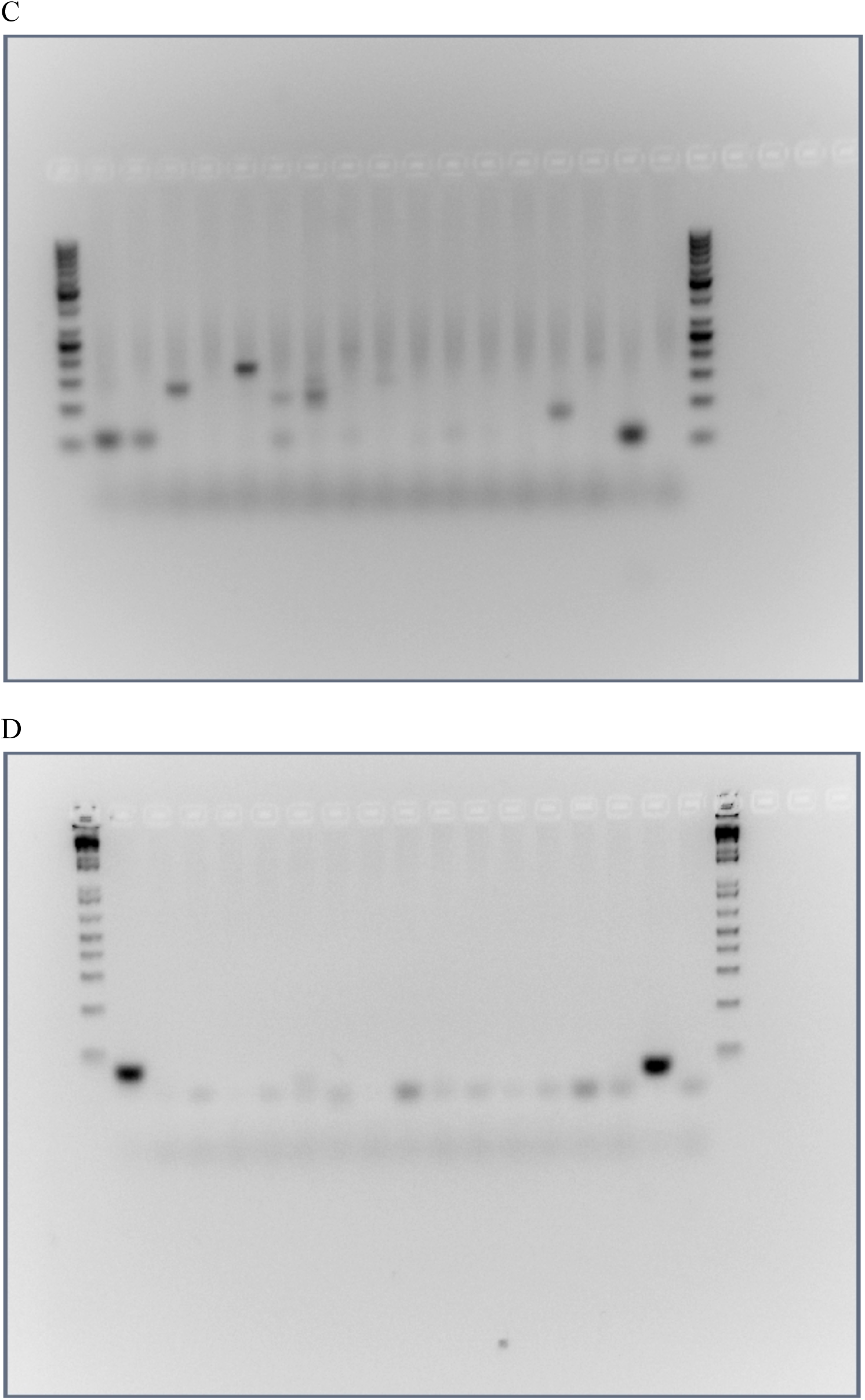
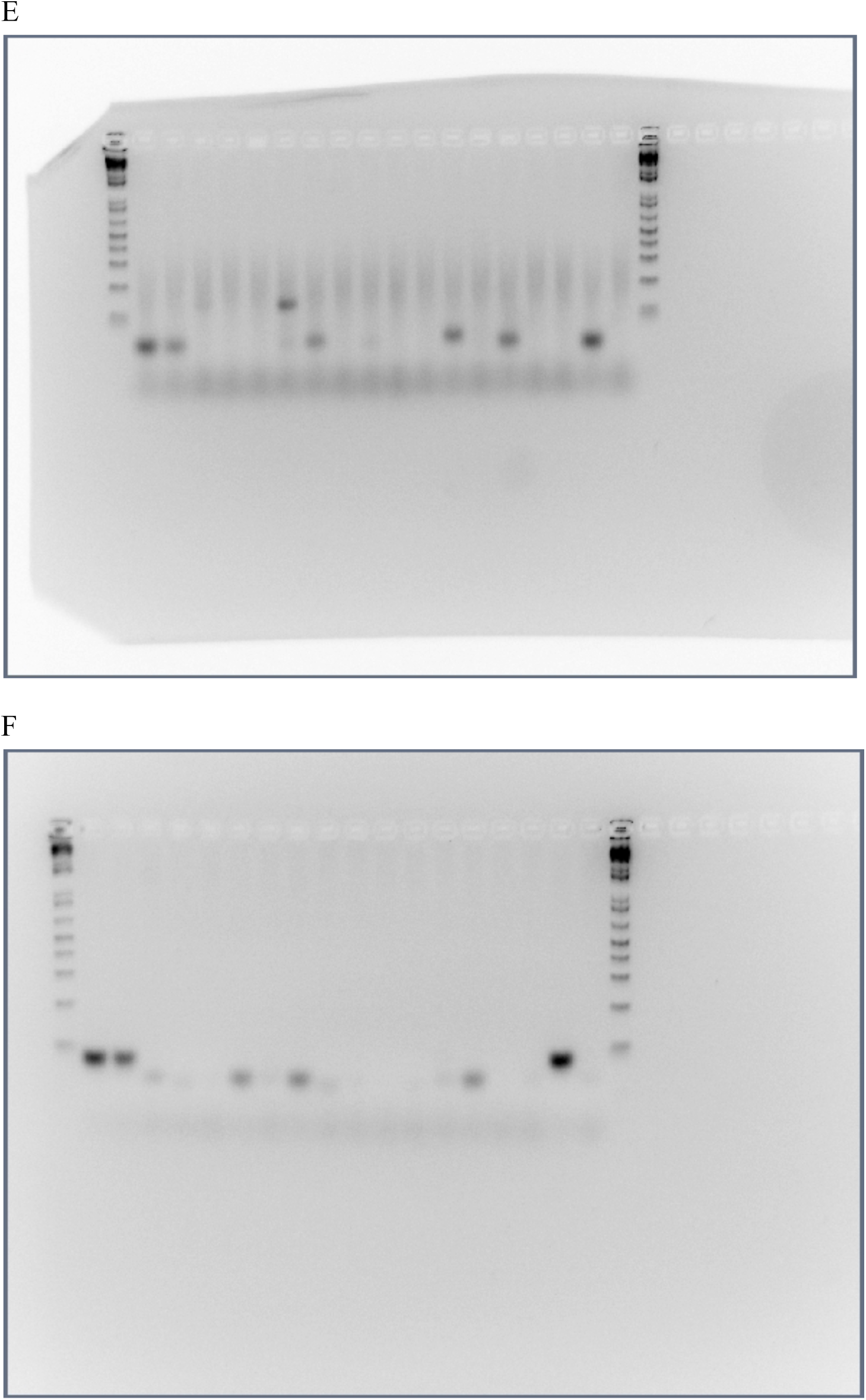
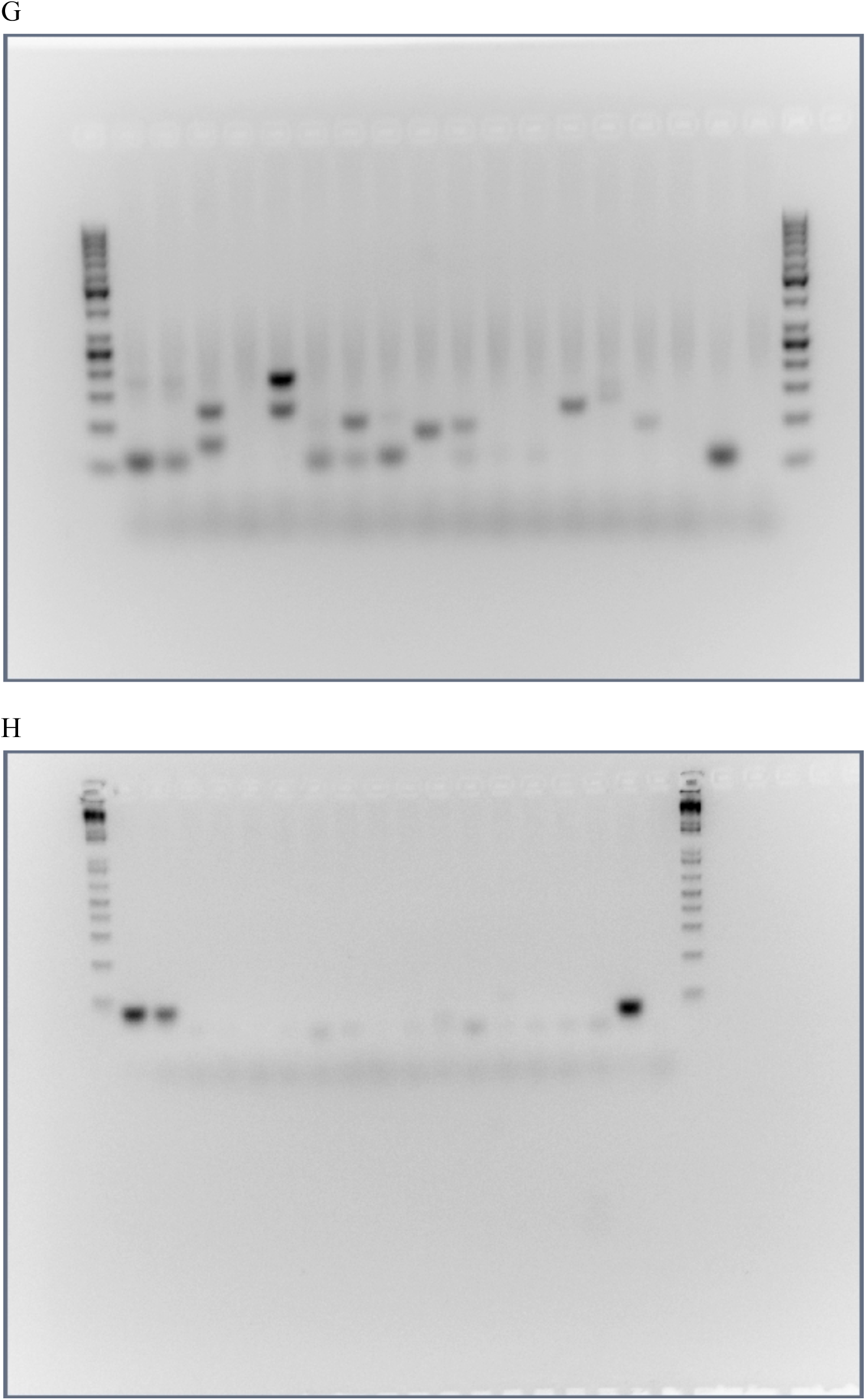
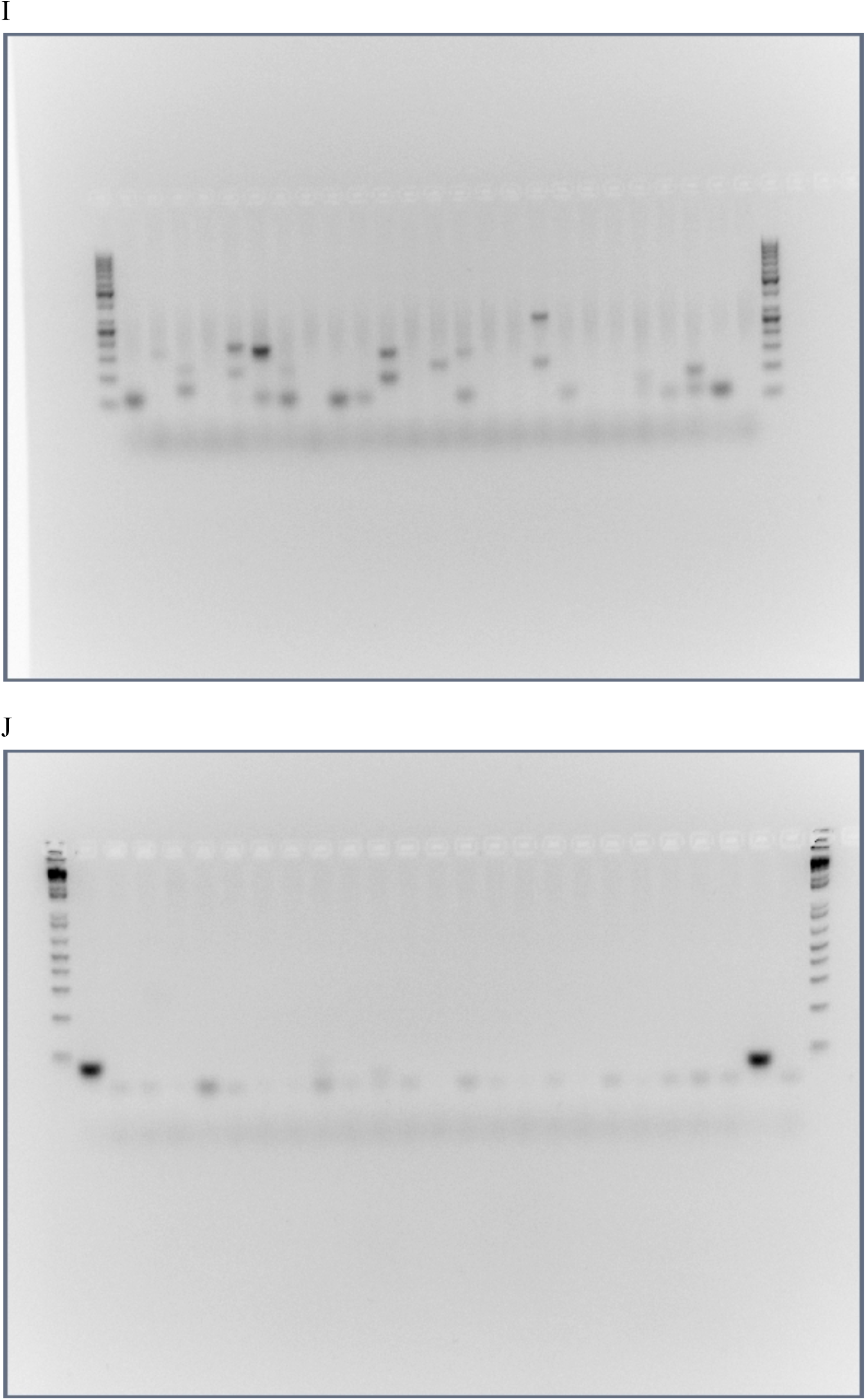
Full size, unedited gels used for Figure 4 in the main text. (**A**) MIR941-Male 1, (**B**) Rabbit β-globin fragment - Male 1, (**C**) MIR941-Male 2, (**D**) Rabbit β-globin fragment - Male 2, (**E**) MIR941-Male 3, (**F**) Rabbit β-globin fragment - Male 3, (**G**) MIR941-Male 4, (**H**) Rabbit β-globin fragment - Male 4, (**I**) MIR941-Male 5, (**J**) Rabbit β-globin fragment - Male 5.

**Supplementary Figure 9.**
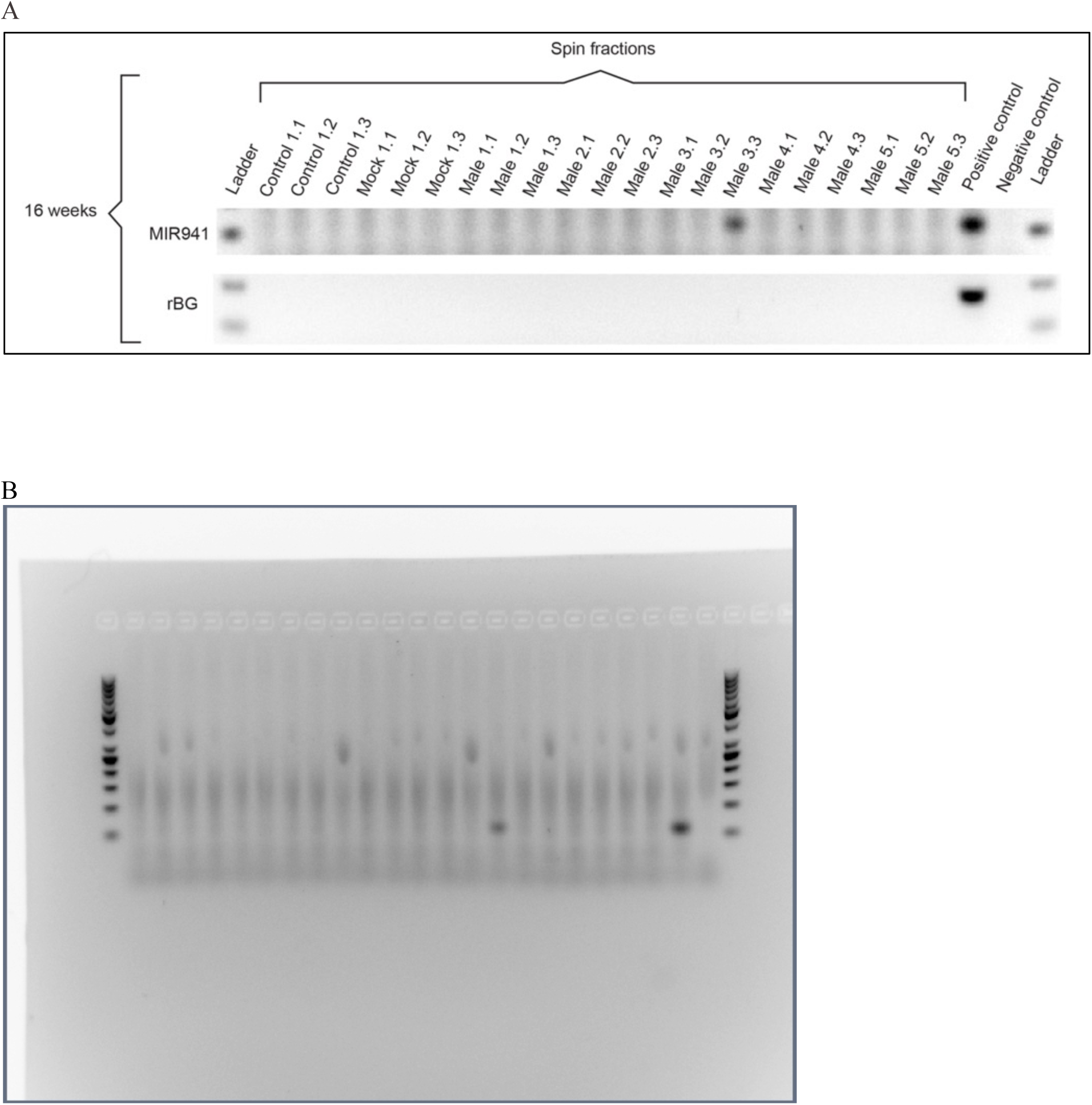
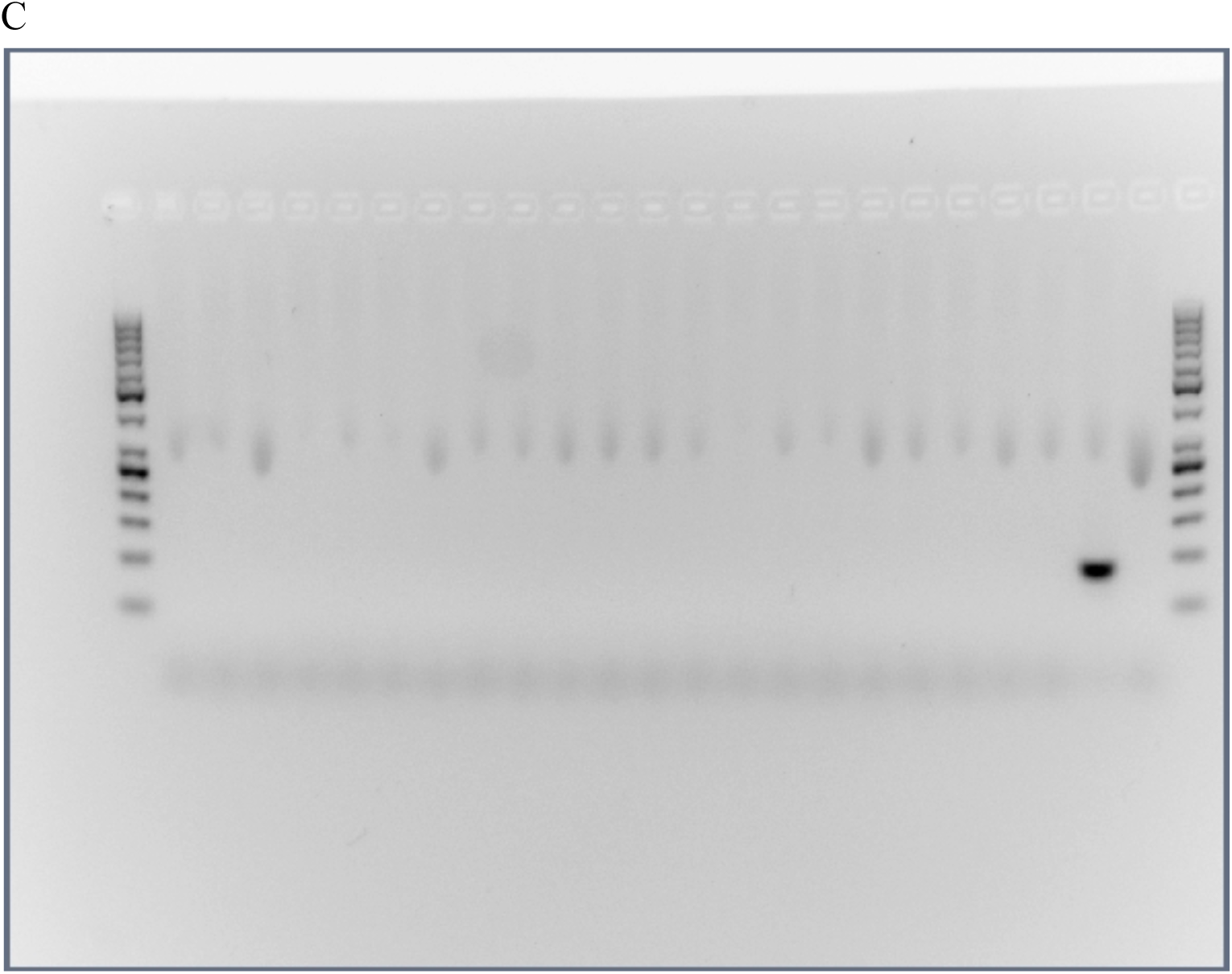
Cropped (**A**) figures showing LNA qPCR products for MIR941 (top panel) and rabbit β-globin fragment (bottom panel) from a series of microcentrifuge and ultracentrifuge spins to isolate various fractions of whole blood in five 16 week treated male mice and one age-matched control and mock-treated male. Spins were (1) 3,000 x g, (2) 12,000 x g and (3) 110,000 x g. Full size, unedited gels are shown in (**B**) MIR941 and (**C**) rabbit β-globin fragment.

## References

1. Carone BR, Fauquier L, Habib N, Shea JM, Hart CE, Li R, Bock C, Li C, Gu H, Zamore PD: Paternally induced transgenerational environmental reprogramming of metabolic gene expression in mammals. Cell 2010, 143: 1084–1096.

2. Yeaman S, Hodgins KA, Lotterhos KE, Suren H, Nadeau S, Degner JC, Nurkowski KA, Smets P, Wang T, Gray LK: Convergent local adaptation to climate in distantly related conifers. Science 2016, 353: 1431–1433.

3. Klosin A, Casas E, Hidalgo-Carcedo C, Vavouri T, Lehner B: Transgenerational transmission of environmental information in C. elegans. Science 2017, 356: 320–323.

4. Singh DP, Saudemont B, Guglielmi G, Arnaiz O, Goût J-F, Prajer M, Potekhin A, Przybòs E, Aubusson-Fleury A, Bhullar S: Genome-defence small RNAs exapted for epigenetic mating-type inheritance. Nature 2014, 509: 447.

5. Öst A, Lempradl A, Casas E, Weigert M, Tiko T, Deniz M, Pantano L, Boenisch U, Itskov PM, Stoeckius M: Paternal diet defines offspring chromatin state and intergenerational obesity. Cell 2014, 159: 1352–1364.

6. Warner DA, Uller T, Shine R: Transgenerational sex determination: the embryonic environment experienced by a male affects offspring sex ratio. Scientific reports 2013, 3: 2709.

7. Bräutigam K, Vining KJ, Lafon-Placette C, Fossdal CG, Mirouze M, Marcos JG, Fluch S, Fraga MF, Guevara MÁ, Abarca D: Epigenetic regulation of adaptive responses of forest tree species to the environment. Ecology and evolution 2013, 3: 399–415.

8. Dias BG, Ressler KJ: Parental olfactory experience influences behavior and neural structure in subsequent generations. Nature neuroscience 2014, 17: 89.

9. Rechavi O, Houri-Ze’evi L, Anava S, Goh WS, Kerk SY, Hannon GJ, Hobert O: Starvationinduced transgenerational inheritance of small RNAs in C. elegans. Cell 2014, 158: 277–287.

10. Bošković A, Rando OJ: Transgenerational epigenetic inheritance. Annual review of genetics 2018, 52: 21–41.

11. Miska EA, Ferguson-Smith AC: Transgenerational inheritance: Models and mechanisms of non-DNA sequence-based inheritance. Science 2016, 354: 59–63.

12. Waldron D: Small RNAs: Regulating transgenerational epigenetics. Nat Rev Genet 2016, 17: 315.

13. Posner R, Toker IA, Antonova O, Star E, Anava S, Azmon E, Hendricks M, Bracha S, Gingold H, Rechavi O: Neuronal Small RNAs Control Behavior Transgenerationally. Cell 2019, 177: 1814–1826 e1815.

14. Houri-Ze’evi L, Korem Y, Sheftel H, Faigenbloom L, Toker IA, Dagan Y, Awad L, Degani L, Alon U, Rechavi O: A Tunable Mechanism Determines the Duration of the Transgenerational Small RNA Inheritance in C. elegans. Cell 2016, 165: 88–99.

15. Lasser C, Eldh M, Lotvall J: Isolation and characterization of RNA-containing exosomes. J Vis Exp 2012: e3037.

16. Cai Q, Qiao L, Wang M, He B, Lin FM, Palmquist J, Huang SD, Jin H: Plants send small RNAs in extracellular vesicles to fungal pathogen to silence virulence genes. Science 2018, 360: 1126–1129.

17. Thomou T, Mori MA, Dreyfuss JM, Konishi M, Sakaguchi M, Wolfrum C, Rao TN, Winnay JN, Garcia-Martin R, Grinspoon SK, et al: Adipose-derived circulating miRNAs regulate gene expression in other tissues. Nature 2017, 542: 450–455.

18. Guduric-Fuchs J, O’Connor A, Camp B, O’Neill CL, Medina RJ, Simpson DA: Selective extracellular vesicle-mediated export of an overlapping set of microRNAs from multiple cell types. BMC Genomics 2012, 13: 357.

19. Skinner MK, Ben Maamar M, Sadler-Riggleman I, Beck D, Nilsson E, McBirney M, Klukovich R, Xie Y, Tang C, Yan W: Alterations in sperm DNA methylation, non-coding RNA and histone retention associate with DDT-induced epigenetic transgenerational inheritance of disease. Epigenetics Chromatin 2018, 11: 8.

20. Ben Maamar M, Sadler-Riggleman I, Beck D, McBirney M, Nilsson E, Klukovich R, Xie Y, Tang C, Yan W, Skinner MK: Alterations in sperm DNA methylation, non-coding RNA expression, and histone retention mediate vinclozolin-induced epigenetic transgenerational inheritance of disease. Environ Epigenet 2018, 4: dvy010.

21. Sarangdhar MA, Chaubey D, Srikakulam N, Pillai B: Parentally inherited long non-coding RNA Cyrano is involved in zebrafish neurodevelopment. Nucleic Acids Res 2018, 46: 9726–9735.

22. An T, Zhang T, Teng F, Zuo JC, Pan YY, Liu YF, Miao JN, Gu YJ, Yu N, Zhao DD, et al: Long non-coding RNAs could act as vectors for paternal heredity of high fat diet-induced obesity. Oncotarget 2017, 8: 47876–47889.

23. Soumillon M, Necsulea A, Weier M, Brawand D, Zhang X, Gu H, Barthes P, Kokkinaki M, Nef S, Gnirke A, et al: Cellular source and mechanisms of high transcriptome complexity in the mammalian testis. Cell Rep 2013, 3: 2179–2190.

24. Hu HY, He L, Fominykh K, Yan Z, Guo S, Zhang X, Taylor MS, Tang L, Li J, Liu J, et al: Evolution of the human-specific microRNA miR-941. Nat Commun 2012, 3: 1145.

25. Foust KD, Nurre E, Montgomery CL, Hernandez A, Chan CM, Kaspar BK: Intravascular AAV9 preferentially targets neonatal neurons and adult astrocytes. Nat Biotechnol 2009, 27: 59–65.

26. Hammond SL, Leek AN, Richman EH, Tjalkens RB: Cellular selectivity of AAV serotypes for gene delivery in neurons and astrocytes by neonatal intracerebroventricular injection. PLoS One 2017, 12: e0188830.

27. Hutson TH, Kathe C, Moon LD: Trans-neuronal transduction of spinal neurons following cortical injection and anterograde axonal transport of a bicistronic AAV1 vector. Gene Ther 2016, 23: 231–236.

28. Xiao L, Bornmann C, Hatstatt-Burkle L, Scheiffele P: Regulation of striatal cells and goaldirected behavior by cerebellar outputs. Nat Commun 2018, 9: 3133.

29. Hong BS, Cho JH, Kim H, Choi EJ, Rho S, Kim J, Kim JH, Choi DS, Kim YK, Hwang D, Gho YS: Colorectal cancer cell-derived microvesicles are enriched in cell cycle-related mRNAs that promote proliferation of endothelial cells. BMC Genomics 2009, 10: 556.

30. Zhou R, Chen KK, Zhang J, Xiao B, Huang Z, Ju C, Sun J, Zhang F, Lv XB, Huang G: The decade of exosomal long RNA species: an emerging cancer antagonist. Mol Cancer 2018, 17: 75.

